# Loss of p27^Kip1^ causes metabolic reprogramming and is sufficient to induce a Warburg effect and glutamine addiction in untransformed cells

**DOI:** 10.64898/2026.02.06.703945

**Authors:** Leo Rolland, Elodie Mitri, Christine Dozier, Marion Aguirrebengoa, Ivan Nemazanyy, Carine Joffre, Jean-Emmanuel Sarry, Anastassia Hatzoglou, Arnaud Besson

**Affiliations:** Molecular, Cellular and Developmental biology department (MCD), Centre de Biologie Integrative (CBI), Université de Toulouse, CNRS, UPS, 31062 Toulouse, France; Platform for Metabolic Analyses, Structure Fédérative de Recherche Necker, INSERM US24/CNRS UAR 3633, Paris, France; Centre de Recherches en Cancérologie de Toulouse, U1037, Inserm, Université de Toulouse, Toulouse, France

## Abstract

The metabolic needs of a cell are tightly linked to its proliferative state and increasing evidence indicate an extensive bidirectional crosstalk between metabolic pathways and cell cycle regulators. In cancer cells, metabolism is reprogrammed to couple energetic needs and relentless proliferation. The cyclin/CDK inhibitor p27^Kip1^ (p27) is frequently inactivated in cancers. p27 is also involved in multiple cellular processes, including transcriptional regulation or autophagy induction. Herein, we investigated the effect of p27 loss on cell metabolism. The knockout of p27 in immortalized mouse fibroblasts increases glucose uptake and glycolysis, while decreasing mitochondrial ATP production, consistent with induction of a Warburg effect, and this was accompanied by an increased glutamine dependency to feed the TCA cycle. Our data suggest that p27 loss causes this phenotype through extensive transcriptional remodeling of metabolic gene expression. Importantly, p27 silencing in human retinal RPE1-hTERT cells was sufficient to induce a Warburg effect. Together, these results reveal a new function of p27 in regulating energy metabolism and that loss of p27 expression is sufficient to induce metabolic reprogramming and a Warburg effect, suggesting that p27 inactivation in cancer cells not only results in the loss of cell cycle inhibition but also enables the metabolic rewiring needed for increased proliferation.

## Introduction

In living organisms, cell division and metabolism must be strictly regulated to couple energetic needs with cell cycle progression. While it was initially thought that the metabolic state of a cell controlled its capacity to divide, increasing evidence now indicates that cell cycle regulators are able to regulate cellular metabolism in a bidirectional crosstalk that allows the fine-tuning of these two essential processes ^1–4^. This ensures that ATP, nutrients and building blocks, such as amino acids and nucleotides, are present in sufficient amounts to carry out the corresponding cell cycle phase. Indeed, CDKs have been found to control metabolism in multiple ways, either by directly phosphorylating metabolic enzymes or metabolic regulators, such as AMPK or mTOR, or indirectly by regulating the activity of transcription factors that control the expression of metabolic enzymes ^1–6^. The CDK inhibitors (CKIs) p21^Cip1^ (p21) or p16^INK4A^ (p16) have also been shown to regulate cell metabolism ^7–9^.

p27^Kip1^ is a cell cycle inhibitor and tumor suppressor via its ability to block cyclin-CDK activity, playing a key role in establishing and maintaining quiescence ^10–12^. Indeed, due to the lack of control over cell proliferation, p27^-/-^ mice are 30% larger than p27^+/+^ animals and spontaneously develop multiple organ hyperplasia and pituitary tumors ^13^. Although the *CDKN1B* gene, encoding p27, is rarely mutated in cancers, p27 is frequently inactivated via either increased proteolytic degradation or exclusion from the nucleus ^10–12,14,15^. p27 also controls other cellular processes, including migration, invasion and transcription and may act as an oncogene ^10,12,16–18^. Several studies have described a role for p27 in the control of autophagy and survival in response to metabolic stress ^19–22^. Depending on the type of nutrient deprivation, p27 regulates autophagy flux by distinct mechanisms. In absence of amino acids, p27 promotes autophagy by participating in the inhibition of mTORC1 by interfering with the assembly of the Ragulator complex on the lysosomal membrane ^20^. In contrast, in absence of glucose, p27 promotes autophagic vesicle trafficking by controlling ATAT1-mediated microtubule acetylation ^19^. p27 is also known to act as a transcriptional co-regulator by interacting with various transcriptional corepressors and transcription factors, via both CDK-dependent and independent mechanisms ^11,23–28^. Interestingly, p27 was found to interact with promoters of genes involved in mitochondrial organization and mitochondrial respiration ^23^. A recent study also described that during caffeine uptake, a fraction of p27 is recruited to the mitochondria where it promotes mitochondrial respiration ^29^. However, while the role of p27 in induction of autophagy under metabolic stress has been rather extensively studied ^19–22^, its potential involvement in the control of energy metabolism in homeostatic conditions remains elusive.

Cellular metabolism includes both energy consuming anabolic or synthesis reactions and catabolic or degradation reactions that produce energy ^30^. Animal cells derive most of their energy production through bioenergetic pathways utilizing glucose, fatty acid or glutamine. Glucose metabolism is initiated via glycolysis, leading to the formation of pyruvate, which joins the tricarboxylic acid (TCA) cycle to provide protons for mitochondrial respiration when oxygen is available. However, if oxygen is limiting, pyruvate is converted into lactate, allowing rapid ATP synthesis when mitochondrial ATP production is impaired^30^. Importantly, most cancer cells exhibit a reprogrammed glucose metabolism, a phenomenon known as the Warburg effect, to support the vast energetic demand required for their relentless cell division ^31^. In the Warburg effect, cells adopt a glycolytic behavior and glucose is preferentially used to generate lactate instead of being used to perform mitochondrial respiration, even in presence of oxygen, leading to a decreased mitochondrial oxygen consumption ^32^. Lactate overproduction is mediated by lactate dehydrogenase (LDH) complexes, which control glucose fate after glycolysis by acting on the balance between pyruvate and lactate ^33–35^. This balance depends on the proportion of the enzymes LDHA and LDHB in the tetrameric LDH complex, which preferentially convert pyruvate into lactate or lactate into pyruvate, respectively ^33–36^. In cancers, LDHA is often upregulated while LDHB is downregulated, leading to lactate overproduction ^37–43^. While glycolysis produces less ATP than complete oxidation of glucose within the mitochondria, an increased glycolytic flux allows generating sufficient ATP levels to maintain cell homeostasis ^44,45^. In cancers, this enhanced glycolytic flux is supported by the overexpression of glucose transporters, such as GLUT1, and of glycolytic enzymes ^46–48^. In addition, the Warburg effect is often associated with increased glutamine uptake and metabolism, through glutaminolysis, providing additional substrate for both mitochondrial respiration and nucleotide biosynthesis ^30,49^. Indeed, the glutamine transporter SLC1A5 and the glutaminases GLS1 and GLS2 are overexpressed in many types of cancers, supporting this glutamine addiction ^50–58^.

In the present study, we investigated how p27 may regulate metabolism using bioenergetics, metabolomics and transcriptomics approaches. We found that the loss of p27 expression is accompanied by an extensive metabolic reprogramming mediated at least in part through transcriptional changes, causing a Warburg effect, coupled with an increased glutamine dependency. These results suggest that loss of p27 expression in cancers could be sufficient to induce a Warburg effect, and illustrate the dual role of p27 in regulating the cell cycle and cell metabolism.

## Results

### p27 loss decreases oxygen consumption rate and increases extracellular acidification

Cells require energy in the form of ATP for growth, proliferation and to maintain homeostatic functions ^59^. ATP is produced through catabolic processes, mostly glycolysis and mitochondrial oxidative phosphorylation (OXPHOS), and scavenging mechanisms such as autophagy ^59^. Since p27 was previously reported to promote autophagy during nutrient starvation and to regulate mTOR activity ^19–21^, we investigated whether p27 could be more generally involved in the regulation of cellular metabolism. For this, the oxygen consumption rate (OCR), as a measure of mitochondrial function, and the extracellular acidification rate (ECAR), which approximates glycolytic activity, were monitored using a Seahorse analyzer in p27^+/+^ and p27^-/-^ immortalized mouse embryonic fibroblasts (MEFs). In these experiments, mitochondrial respiration can be dissected using drugs affecting the activity of the mitochondrial respiratory chain or mitochondrial membrane potential (Figure 1a, left panel). Strikingly, OCR was reduced in p27^-/-^ cells compared to p27^+/+^ (Figure 1a, right panel), with a significant decrease of mitochondrial basal respiration (Figure 1b), maximal respiratory capacity (Figure 1c) and spare capacity (Figure 1d). Importantly, ATP-linked OCR, indicative of the oxygen level consumed by mitochondria to generate ATP, was significantly decreased in p27^-/-^ MEFs (Figure 1e). On the other hand, non-mitochondrial OCR (Figure 1f) and proton leak (Figure 1g) were not significantly affected by p27 status, confirming that decreased OCR in p27^-/-^ cells is a consequence of reduced mitochondrial oxygen consumption. Flow cytometry analyses of cells stained with Mitotracker Green FM (MTG) to determine mitochondrial mass did not reveal any significant difference between p27^+/+^ and p27^-/-^ MEFs (Figure 1h). Together, these results indicate that mitochondrial respiration is decreased in absence of p27 expression.

**Figure 1:**
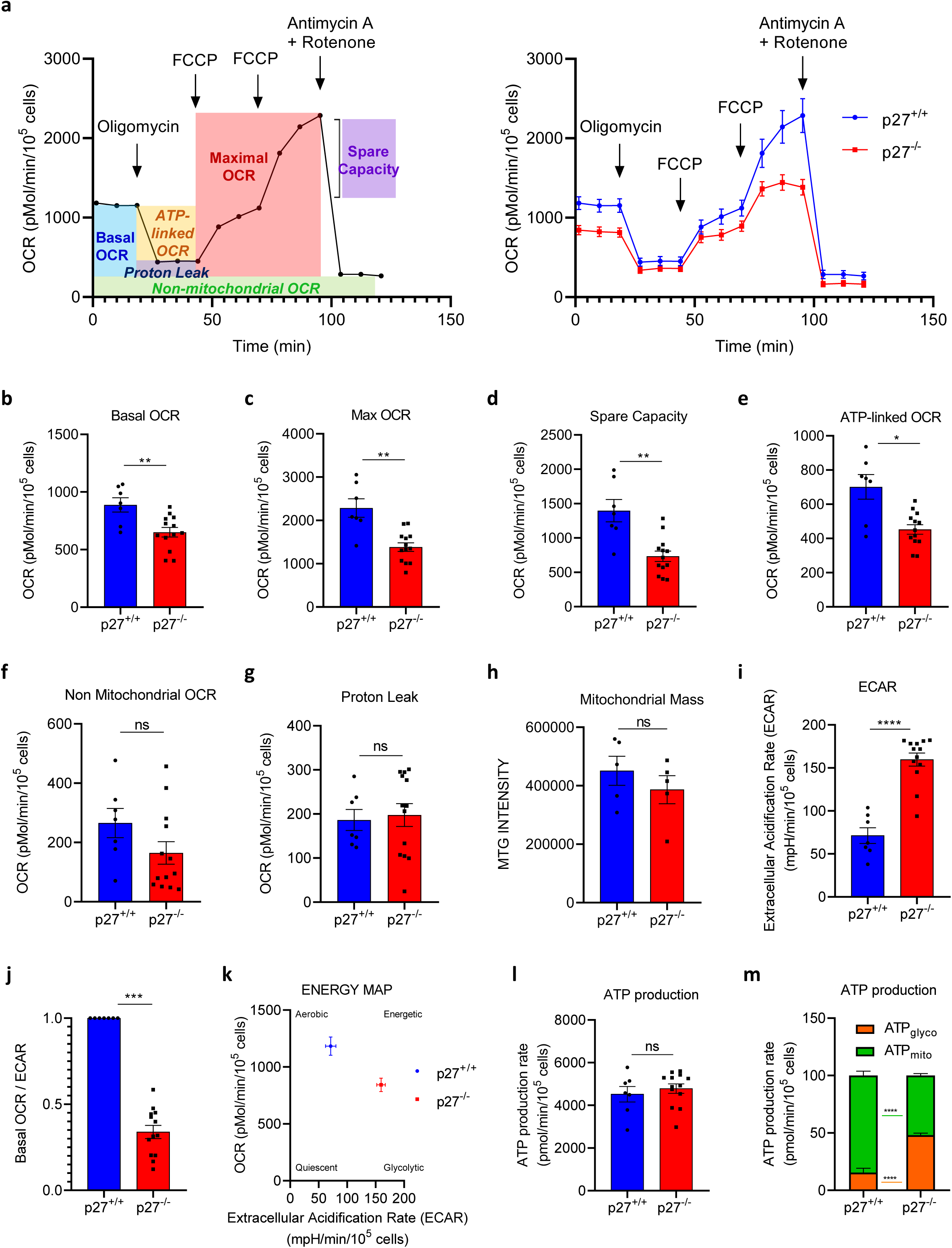
p27 loss decreases OCR and increases ECAR. **(a)** Oxygen consumption rate (OCR) was measured at various time points with consecutive injections of oligomycin (1 µM), FCCP (two injections at 2 µM and 6 µM), and Rotenone/antimycin A (1 µM each), and normalized to cell number. The left panel shows a representative OCR curve with the calculations used to determine the different components of OCR in Seahorse Mitostress experiments. The right panel shows the mean OCR curves of p27^+/+^ (n = 7) and p27^-/-^ (n = 13) MEFs. **(b-g)** Basal respiration (b), Maximal respiration (c), Spare capacity (d), ATP-linked respiration (e), Non-mitochondrial OCR (f) and Proton leak (g) of p27^+/+^ (n = 7) and p27^-/-^ (n = 13) MEFs, derived from Seahorse Mitostress experiments (a, right panel). Graphs show means ± SEM. ns: p > 0.05; *: p < 0.05; **: p < 0.01. **(h)** Mitochondrial Mass was determined by flow cytometry analysis of Mitotracker Green FM (MTG)-stained p27^+/+^ and p27^-/-^ MEFs. Graph shows mean MTG intensity ± SEM from 5 independent experiments. ns: p > 0.05. **(i)** Extracellular acidification rate (ECAR) of p27^+/+^ (n = 7) and p27^-/-^ (n = 13) MEFs. Graph shows means ± SEM. ****: p < 0.0001. **(j)** OCR / ECAR ratio of p27^+/+^ (n = 7) and p27^-/-^ (n = 13) MEFs from Seahorse Mitostress experiments. Graphs show means ± SEM. ***: p < 0.001. **(k)** Energy map of p27^+/+^ (n = 7) and p27^-/-^ (n = 13) MEFs from Seahorse Mitostress experiments. **(l, m)** ATP production in p27^+/+^ (n = 7) and p27^-/-^ (n = 13) MEFs (k), and Glycolytic ATP/Mitochondrial ATP production ratio (l) based on Seahorse Real-time ATP rate Assays. Graphs show means ± SEM. ns: p > 0.05; ****: p < 0.0001.

Although less energetically favorable than OXPHOS, glycolysis is the other main ATP-generation mechanism in the cell, and is favored in cancer cells and known as the Warburg effect ^59^. Since ATP-linked OCR is decreased in p27^-/-^ cells, we asked whether this was accompanied by an increase in ECAR, which reflects lactate production via glycolysis. ECAR was dramatically increased in p27^-/-^ cells (Figure 1i), suggesting an increased glycolytic flux. Consistently, the basal OCR/ECAR ratio was decreased in p27^-/-^ cells (Figure 1j), suggesting that p27 loss drives metabolism toward a glycolytic behavior and modifies the glycolytic/mitochondrial ATP production balance. To appreciate the effect of p27 on the metabolic state of the cell, an energy map was created using OCR and ECAR data, clearly indicating that p27 loss shifts the metabolic state from an aerobic to a glycolytic behavior (Figure 1k). To further dissect the role of p27, the glycolytic and mitochondrial ATP production were estimated by combining OCR and ECAR data from the Seahorse analyzer ^60^. While total ATP production was similar between p27^+/+^ and p27^-/-^ MEFs (Figure 1l), p27^-/-^ cells produced more ATP via glycolysis (48.1%) whereas p27^+/+^ cells produced their ATP mostly via mitochondrial respiration (84.6%) (Figure 1m). In summary, our results suggest that p27 promotes ATP production via mitochondrial respiration, while the loss of p27 induces a rewiring of metabolic pathways leading to ATP production via glycolysis.

### p27 loss enhances glycolysis and promotes the Warburg effect

To explore how p27 loss enhances glycolysis, Seahorse glycolysis stress experiments were performed to evaluate the different components of ECAR (Figure 2a, left panel). Under these conditions, cells were initially grown in absence of glucose for 1 h and then glucose, Oligomycin and 2-deoxy-D-glucose (2-DG) were added successively to determine glycolytic and non-glycolytic ECAR. Consistent with our previous observation (Figure 1i), there was an overall increase of ECAR in p27^-/-^ cells compared to p27^+/+^ MEFs (Figure 2a, right panel). Surprisingly, basal ECAR, reflecting lactate production in absence of glucose, was significantly higher in p27^-/-^ MEFs (Figure 2b). One possible explanation is that a short-term glucose starvation (1 h) is not sufficient to fully deplete intracellular glucose in these cells. Upon glucose addition, glycolysis-linked ECAR was markedly increased in p27^-/-^ cells, indicating that p27 promotes glycolytic flux when glucose is available (Figure 2c). Next, the glycolytic capacity, measured after ATP synthase inhibition with Oligomycin, was also increased in p27^-/-^ MEFs, further supporting that p27 promotes glycolysis to generate ATP even when mitochondrial activity is impaired (Figure 2d). The glycolytic reserve was independent of p27 status (Figure 2e), suggesting that p27 does not affect the ability to rapidly shift toward a glycolytic behavior when mitochondrial activity is impaired. Importantly, there was no difference in non-glycolytic acidification between p27^+/+^ and p27^-/-^ MEFs (Figure 2f), further confirming that the increased ECAR in p27^-/-^ cells is due to increased glucose metabolism, either via increased glycolysis or increased pyruvate conversion into lactate, or both.

**Figure 2:**
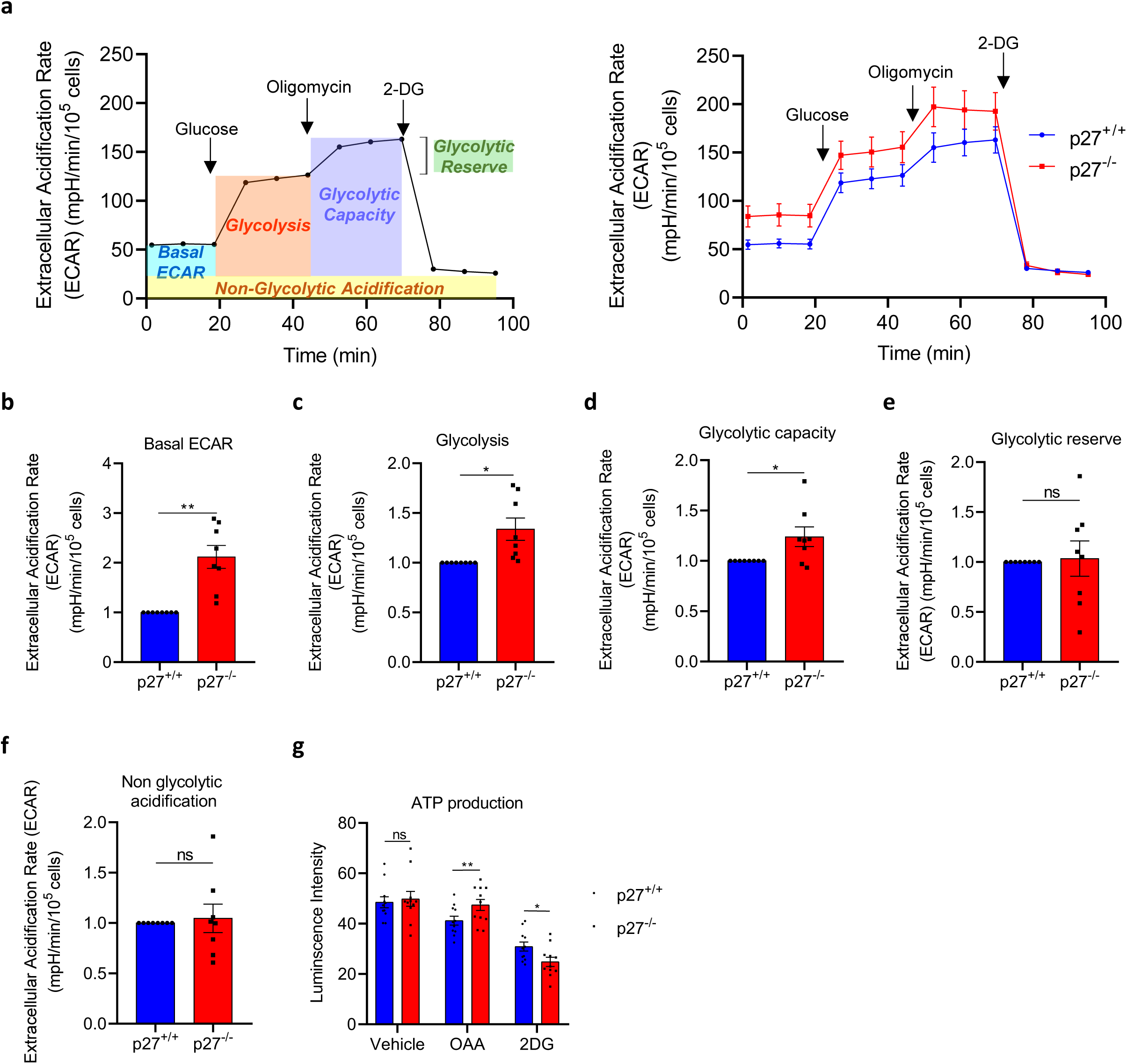
p27 loss confers a glycolytic behavior, causing a Warburg effect. **(a)** Extracellular acidification rate (ECAR) was measured at various time points with consecutive injections of glucose (10 mM), oligomycin (1 μM), and 2-DG (50 mM) and normalized to cell number. The left panel shows a representative ECAR curve with the calculations used to determine the different components of ECAR in Seahorse Glycolysis Stress experiments. The right panel shows the mean ECAR curves of p27^+/+^ (n = 8) and p27^-/-^ (n = 8) MEFs. **(b-f)** Basal ECAR (b), Glycolysis (c), Glycolytic capacity (d), Glycolytic reserve (e) and Non-glycolytic acidification (f) of p27^+/+^ (n = 8) and p27^-/-^ (n = 8) MEFs, derived from Seahorse Glycolysis Stress experiments (a, right panel). Graphs show means ± SEM. ns: p > 0.05; *: p < 0.05; **: p < 0.01. **(g)** ATP quantification using Firefly Luciferase ATP Assay kit of p27^+/+^ and p27^-/-^ MEFs (n = 11) treated with either vehicle, Oligomycin + Antimycin A (OAA) (10 µM each) or 2-DG (50 mM). Graph shows means ± SEM. ns: p > 0.05; *: p < 0.05; **: p < 0.01.

To confirm the observations obtained with the Seahorse analyzer, ATP was quantified in p27^+/+^ and p27^-/-^ MEFs using Firefly Luciferase ATP assays. There was no significant difference in total ATP production between p27^+/+^ and p27^-/-^ (Figure 2g), in agreement with the estimation of ATP production from Seahorse data (Figure 1l). Then, to determine the importance of the different pathways leading to ATP production, p27^+/+^ and p27^-/-^ MEFs were treated with Oligomycin A and Antimycine A (OAA) to bloc mitochondria-linked ATP production or 2-DG to bloc glycolysis-linked ATP production. When mitochondrial ATP production was blocked, p27^+/+^ cells produced less ATP, while p27^-/-^ cells were unaffected (Figure 2g), consistent with the idea that p27^+/+^ MEFs produce most of their ATP through mitochondrial OXPHOS (Figure 1m). On the contrary, p27^-/-^ MEFs were more sensitive to inhibition of ATP production through glycolysis than p27^+/+^ cells (Figure 2g), confirming that p27^-/-^ cells adopt a glycolytic behavior for ATP production. Taken together, our data shows that p27 loss induces a metabolic shift from mitochondrial ATP production to a glycolytic ATP production, as described in the Warburg effect.

### p27 loss affects global cell metabolism in both normal and metabolic stress conditions

To better understand the role of p27 in the regulation of bioenergetic pathways, we performed targeted metabolic profiling using mass spectrometry. Metabolome (cells metabolites) and exometabolome (metabolites secreted or consumed in culture media) were analyzed in cells grown for 24 h in either complete medium, or in glucose or amino acid starvation medium. Volcano plots (Figure 3, left panels) represent the set of statistically different metabolites and exometabolites between p27^+/+^ and p27^-/-^MEFs, which were used to generate heatmaps (Figure 3, right panels). Dendrograms and Principal Component Analysis representing the metabolite variability for each cell culture condition shows the clustering of cells according to their genotype (Figure S1).

**Figure 3:**
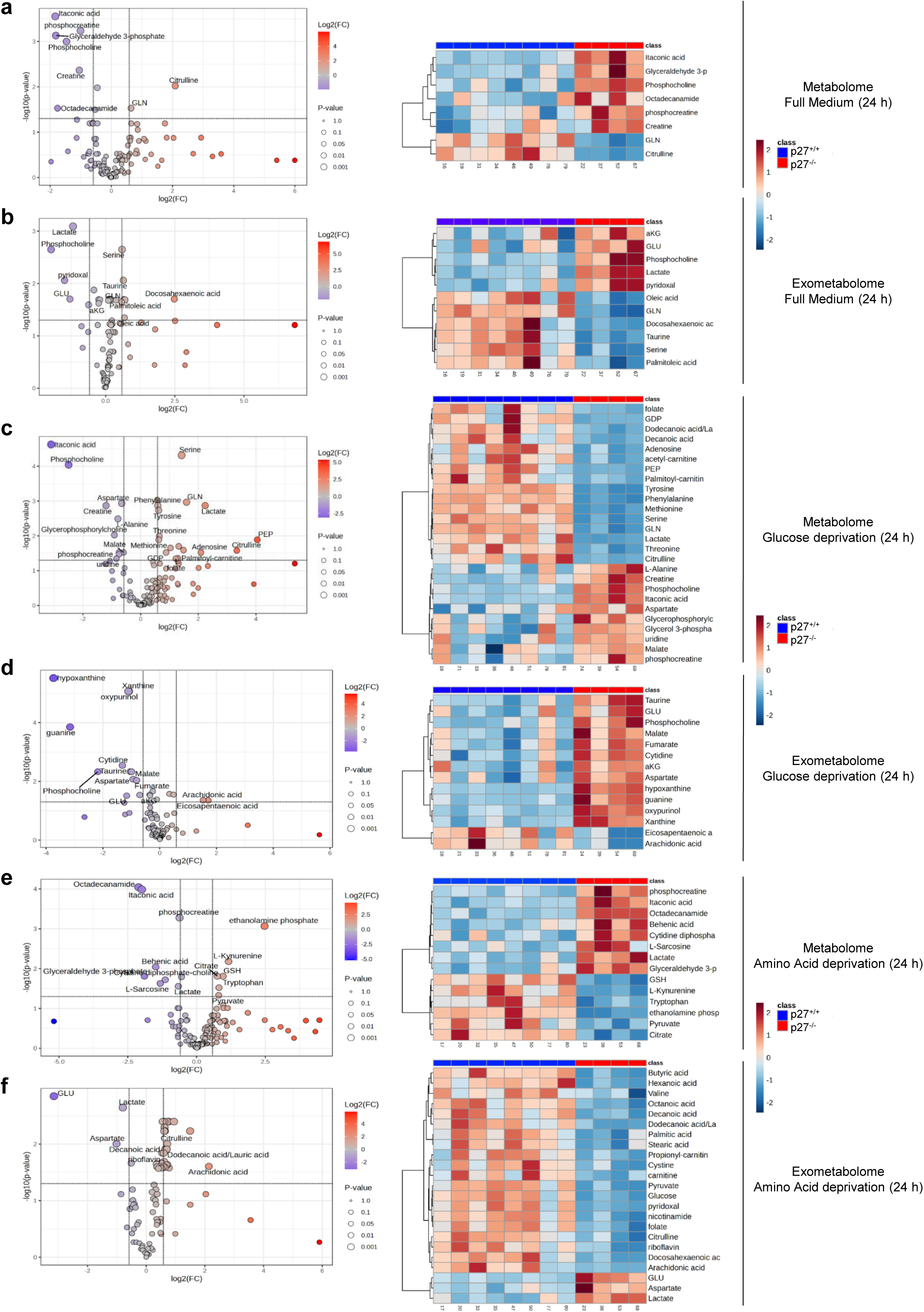
p27 loss affects global cell metabolism in both normal and metabolic stress conditions. **(a, b)** Quantitative analyses of metabolic alterations between p27^+/+^ (n = 8) and p27^-/-^ (n = 4) MEFs (metabolome) (a) or in the corresponding culture media (exometabolome) (b) of cells grown for 24 h in full medium. In the volcano plots (left panels), all the metabolites significantly altered [log2(Fold Change) > 0.585 or Log2(FC) < −0.585, corresponding to a fold change of ± 1.5 x; and False Discovery Rate (FDR) < 0.05] between p27^+/+^ and p27^-/-^ MEFs are named. The heatmaps (left panels) show only the metabolites (a) or exometabolites (b) significantly altered between p27^+/+^ and p27^-/-^ MEFs. **(c-f)** Similar experiments were performed on p27^+/+^ (n = 8) and p27^-/-^ (n = 4) MEFs grown for 24 h in either glucose starvation medium (c, d) or in amino acid starvation medium (e, f).

The absence of p27 modifies the metabolome and exometabolome of cells grown in full medium (Figure 3a, b), confirming that p27 controls cellular metabolism under homeostatic conditions. Interestingly, glyceraldehyde-3-phosphate (G3P), a glycolysis metabolite, accumulated in p27^-/-^ cells in full medium conditions (Figure 3a), supporting a role for p27 in modulating glycolytic flux. In addition, glutamine, which can enter the TCA cycle ^61^, was decreased in both p27^-/-^ cells and medium, suggesting that glutamine uptake and metabolism may be enhanced in p27^-/-^ MEFs (Figure 3a, b). The decrease of oleic, docosahexaenoic and palmitoleic acids in p27^-/-^ cells suggests a role of p27 in the regulation of fatty acid metabolism (Figure 3b). Importantly, lactate levels were markedly higher in p27^-/-^ medium, consistent with the increased ECAR and glycolytic behavior observed in p27^-/-^ cells compared to p27^+/+^ MEFs (Figure 3b and Figure 1i).

Under glucose starvation conditions, several amino acids were decreased in p27^-/-^ MEFs (Figure 3c), likely reflecting the defective autophagy previously reported in p27^-/-^ cells ^19^. Possibly due to this decreased autophagy flux, the levels of phosphoenolpyruvate (PEP), a glycolysis metabolite, as well as lactate were reduced in p27^-/-^ MEFs under glucose deprivation conditions, while malate, a TCA cycle metabolite, was increased (Figure 3c), reinforcing the idea that p27^-/-^ cells preferentially use glucose to produce lactate after glycolysis instead of feeding the TCA cycle. Surprisingly, malate and fumarate, both intracellular metabolites, accumulated in p27^-/-^ cell medium (Figure 3d), likely a consequence of the increased sensitivity of p27^-/-^ cells to apoptosis during glucose starvation ^19,20^. As seen in full medium conditions, glutamine was less abundant in p27^-/-^ cells under glucose deprivation (Figure 3c), suggesting that glutamine uptake and metabolism are enhanced in p27^-/-^ cells independently of glucose availability.

During amino acids deprivation, p27 also affected cellular metabolism (Figure 3e, f). First, p27^-/-^ cells exhibited lower levels of glutathione (GSH) (Figure 3e), an antioxidant mediating the disposal of reactive oxygen species (ROS) ^62^, suggesting a role for p27 in ROS metabolism during metabolic stress. Importantly, G3P and lactate levels were elevated in p27^-/-^ compared to p27^+/+^ cells, while pyruvate level was decreased (Figure 3e), confirming that p27 loss drives glycolysis towards lactate production instead of using pyruvate to feed the TCA cycle, even in absence of amino acids. Moreover, many fatty acids (butyric, hexanoic, octanoic, decanoic, dodecanoic, stearic, docosahexaenoic and arachidonic acids) were decreased in p27^-/-^ cells media (Figure 3f), suggesting that they may be used to feed the TCA cycle ^63^ in absence of amino acids, particularly glutamine. Furthermore, glucose levels were less abundant in p27^-/-^cell media (Figure 3f), suggesting that glucose uptake and/or metabolism is enhanced in p27^-/-^ cells during amino acids starvation. In conclusion, these metabolomics studies reveal that p27 is involved in the regulation of cellular metabolism both in homeostatic and metabolic stress conditions and confirm that in absence of p27, cells adopt a glycolytic behavior and suggest that the TCA cycle could be fed by other energy sources such as glutamine or fatty acids.

### Loss of p27 induces the expression of glycolytic enzymes

Previous reports have described a role for p27 as a transcriptional co-regulator ^11,23–28^. To determine whether p27 may regulate bioenergetic metabolism by modulating gene expression, RNA-Seq analyses were performed on p27^+/+^ and p27^-/-^ MEFs grown for 24 h in full medium for bulk RNA profile comparisons. In the entire genome, 15207 genes were expressed at appreciable levels (> 50 reads per kilobase per million mapped reads [RPKM]) in the 12 samples (p27^+/+^ [n = 8] and p27^-/-^ [n = 4]). Genome viewer captures confirmed that there was no p27 coding sequences in p27^-/-^ cells (Figure 4a). Differential DESeq2 analyses revealed that 2407 genes were significantly deregulated in p27^-/-^ MEFs compared to p27^+/+^ cells (i.e. with −0.6 < log2(FC) < 0.6 and p < 0.05), with 1735 genes downregulated and 672 genes upregulated in p27^-/-^ MEFs (Figure 4b).

**Figure 4:**
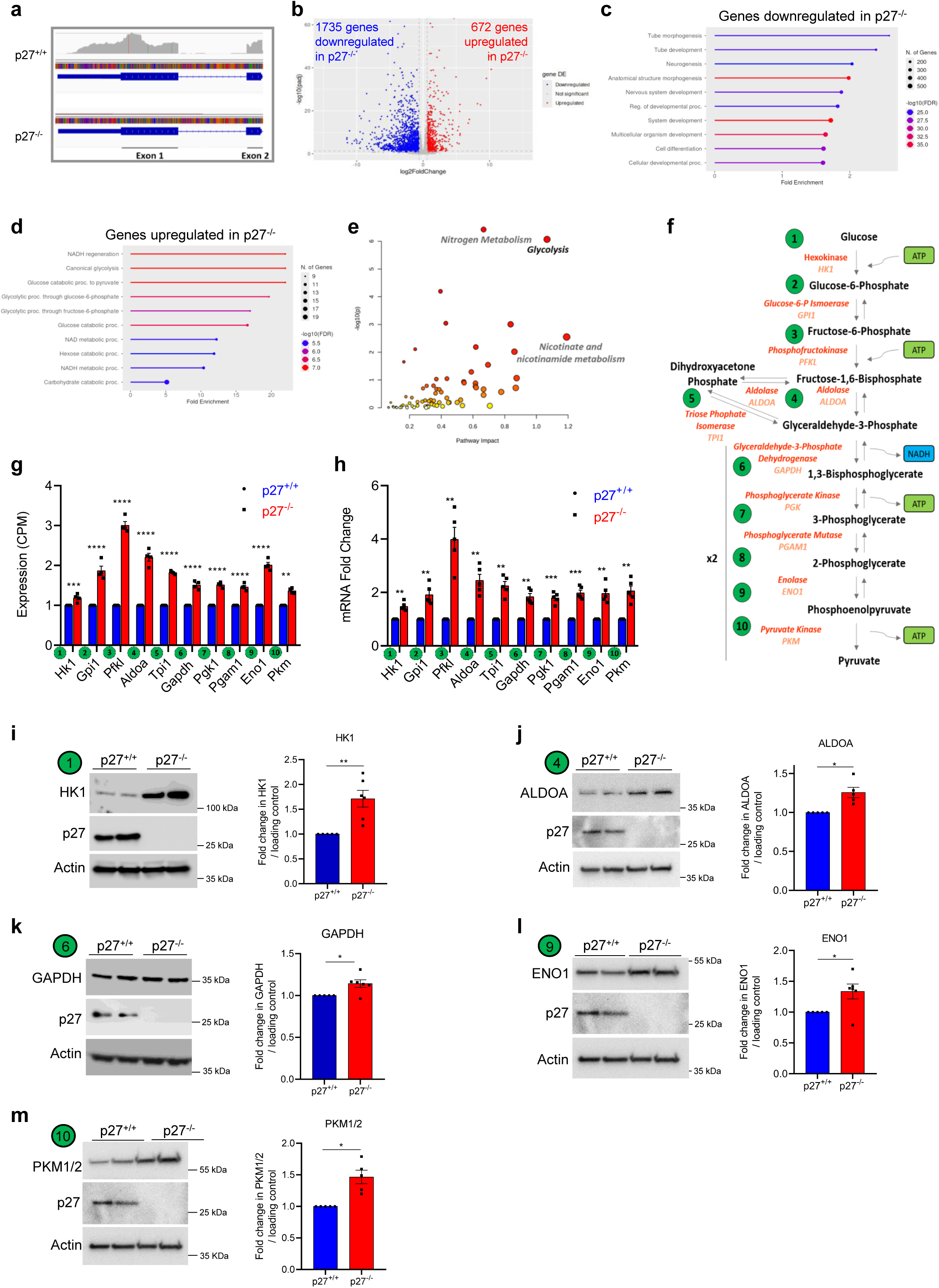
p27 loss induces an upregulation of glycolytic enzymes. **(a)** Example of genome viewer capture of the p27 gene locus from transcriptomics data of p27^+/+^ and p27^-/-^ MEFs. **(b)** Volcano plot showing genes differentially expressed between p27^+/+^ (n = 8) and p27^-/-^ (n = 4) MEFs. Genes were considered differentially expressed when −0.6 < log2(FC) < 0.6 and p < 0.05. **(c, d)** Gene ontology analyses of RNA-Seq data from p27^+/+^ and p27^-/-^ MEFs cultivated in full medium for 24 h. The pathways most affected are shown for genes downregulated (c) and upregulated (d) in p27^-/-^ MEFs. **(e)** Data integration of transcriptomics and metabolomics datasets showing the ontology pathways most deregulated between p27^+/+^ and p27^-/-^ MEFs cultivated in full medium for 24 h. **(f)** Graphical representation of glycolysis showing metabolites (black), enzymes (red) and corresponding genes (orange). **(g)** Gene expression of the 10 glycolytic enzymes in p27^+/+^ (n = 8) and p27^-/-^ (n = 4) MEFs from RNA-Seq data expressed as Count Per Million (CPM), normalized to p27^+/+^ levels. Graph shows means ± SEM. **: p < 0.01; ***: p < 0.001; ****: p < 0.0001. **(h)** RT-qPCR validation of glycolytic enzymes expression in p27^+/+^ and p27^-/-^ MEFs. mRNA levels were normalized to p27^+/+^. Graph shows means ± SEM from five independent experiments. **: p < 0.01; ***: p < 0.001. **(i)** Immunoblots for Hexokinase 1 (HK1), p27 and actin (loading control) in p27^+/+^ and p27^-/-^ MEFs (left panel). Graph shows means ± SEM of the quantification of HK1 levels normalized to actin (p27^+/+^ n = 5; p27^-/-^ n = 6) (right panel). **: p < 0.01. **(j)** Immunoblots for Aldolase (ALDOA), p27 and actin in p27^+/+^ and p27^-/-^ MEFs (left panel). Graph shows means ± SEM of the quantification of ALDOA levels normalized to actin (n = 5) (right panel). *: p < 0.05. **(k)** Immunoblots for Glyceraldehyde-3-Phosphate Dehydrogenase (GAPDH), p27 and actin in p27^+/+^ and p27^-/-^ MEFs (left panel). Graph shows means ± SEM of the quantification of GAPDH levels normalized to actin (p27^+/+^ n = 5; p27^-/-^ n = 6) (right panel). *: p < 0.05. **(l)** Immunoblots for Enolase 1 (ENO1), p27 and actin in p27^+/+^ and p27^-/-^ MEFs (left panel). Graph shows means ± SEM of the quantification of ENO1 levels normalized to actin (p27^+/+^ n = 5; p27^-/-^ n = 6) (right panel). *: p < 0.05. **(m)** Immunoblots for Pyruvate Kinase (PKM1/2), p27 and actin in p27^+/+^ and p27^-/-^ MEFs (left panel). Graph shows means ± SEM of the quantification of PKM1/2 levels normalized to actin (n = 5) (right panel). *: p < 0.05.

Gene ontology analyses of genes downregulated in p27^-/-^ MEFs showed clusters involved in developmental processes, neurogenesis and differentiation (Figure 4c), consistent with previously reported functions of p27 ^23–28^. On the other hand, gene ontology analyses of genes upregulated in p27^-/-^cells revealed an upregulation of genes involved in metabolism, especially glycolysis and carbohydrate metabolism, as well as NAD/NADH metabolism and regeneration (Figure 4d). These analyses support the idea that p27 may control glycolysis by modulating gene expression of glycolytic enzymes. To refine these analyses and identify functional clusters involved in metabolism most enriched in p27^-/-^ MEFs, we performed an integration of RNA-Seq datasets with the metabolomics and exometabolomics datasets described above (Figure 4e). This data integration confirmed that the pathways most deregulated in cells lacking p27 are linked to glycolysis, nicotinamide metabolism and nitrogen metabolism (Figure 4e).

Glycolysis is composed of ten enzymatic reactions resulting in the transformation of glucose into pyruvate (Figure 4f). Strikingly, extraction of the gene expression levels of the ten enzymes responsible for the glycolysis reactions from RNA-Seq data revealed that all of them were significantly upregulated in p27^-/-^ MEFs compared to p27^+/+^ (Figure 4g), which was confirmed by RT-qPCR (Figure 4h). The upregulation of select glycolytic enzymes, namely HK1 (Figure 4i), ALDOA (Figure 4j), GAPDH (Figure 4k), ENO1 (Figure 4l), and PKM1/2 (Figure 4m) in p27^-/-^ MEFs was also confirmed by immunoblot. Altogether, this data confirms the role of p27 as a transcriptional co-regulator and suggest that it actively participates in modulating the expression of genes involved in metabolism and notably in glycolysis.

### p27 loss controls pyruvate fate, leading to lactate overproduction

Pyruvate, the final product of glycolysis, is converted either in Acetyl-coA to enter the TCA cycle or into lactate. Four enzymes control pyruvate fate: pyruvate dehydrogenase (PDH) and its inhibitor pyruvate dehydrogenase kinase 1 (PDK1) control the conversion of pyruvate to Acetyl-coA, while lactate dehydrogenase A/B (LDHA/B) control the pyruvate/lactate balance (Figure 5a) ^64^. Loss of p27 leads to increased ECAR (Figure 1i), suggesting lactate accumulation, which was confirmed by metabolomics studies (Figure 3b and Figure 5b) and this difference was abolished under glucose starvation conditions (Figure 5c), indicating that in absence of p27, glucose metabolism is modified. LDHA was upregulated in p27^-/-^ cells at the mRNA (RNA-Seq and RT-qPCR, Figure 5d, e) and protein levels (Figure 5f) compared to p27^+/+^, while LDHB was downregulated in p27^-/-^ cells compared to p27^+/+^ (Figure 5g-i). In addition, PDK1 was markedly upregulated (Figure 5j-l), suggesting that the conversion of pyruvate to Acetyl-coA is inhibited in p27^-/-^ cells. Overall, these results suggest that the pyruvate/lactate balance is affected in absence of p27, promoting the conversion of pyruvate into lactate instead of directing pyruvate towards the TCA cycle.

**Figure 5:**
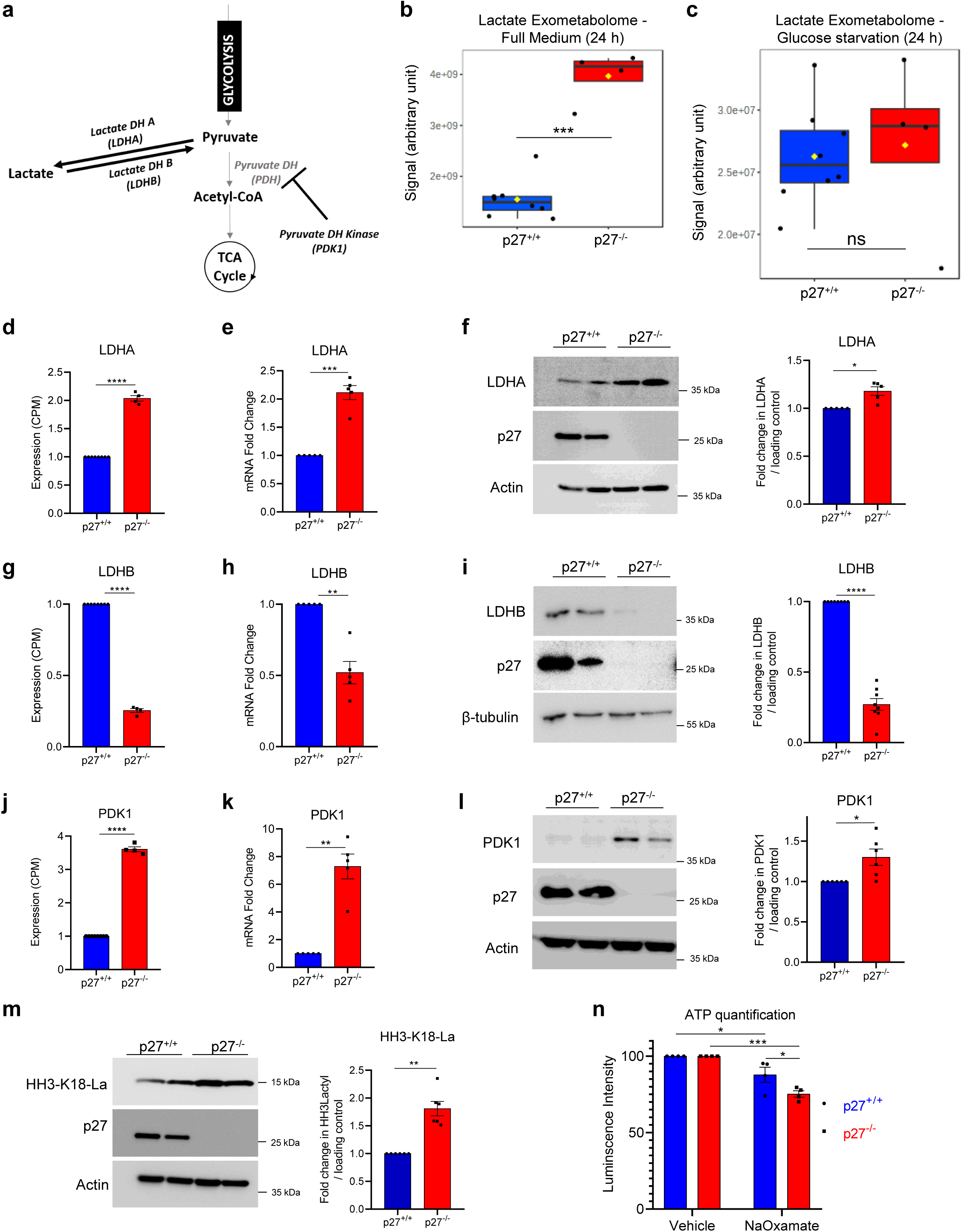
p27 loss controls pyruvate fate, leading to lactate overproduction. **(a)** Graphical representation of pyruvate fate after glycolysis. **(b, c)** Lactate levels in the exometabolome of p27^+/+^ (n = 8) and p27^-/-^ (n = 4) MEFs grown in full medium (b) or glucose starvation medium (c) for 24 h. Data is presented as box plot, means are represented by yellow squares. ns: p > 0.05; ***: p < 0.001. **(d)** LDHA gene expression in p27^+/+^ (n = 8) and p27^-/-^ (n = 4) MEFs grown in full medium for 24 h from RNA-Seq data expressed as Count Per Million, normalized to p27^+/+^ levels. Graph shows means ± SEM. ****: p < 0.0001. (**e**) LDHA mRNA levels in p27^+/+^ and p27^-/-^ MEFs by RT-qPCR, normalized to p27^+/+^ levels. Graph shows means ± SEM from five independent experiments. ***: p < 0.001. (**f**) Immunoblots for LDHA, p27 and actin in p27^+/+^ and p27^-/-^ MEFs (left panel). Graph shows means ± SEM of the quantification of LDHA levels normalized to actin (n = 5) (right panel). *: p < 0.05. (**g**) LDHB gene expression in p27^+/+^ (n = 8) and p27^-/-^ (n = 4) MEFs grown in full medium for 24 h from RNA-Seq data expressed as Count Per Million, normalized to p27^+/+^ levels. Graph shows means ± SEM. ****: p < 0.0001. (**h**) LDHB mRNA levels in p27^+/+^ and p27^-/-^ MEFs by RT-qPCR, normalized to p27^+/+^ levels. Graph shows means ± SEM from five independent experiments. **: p < 0.01. (**i**) Immunoblots for LDHB, p27, and β-tubulin in p27^+/+^ and p27^-/-^ MEFs (left panel). Graph shows means ± SEM of the quantification of LDHB levels normalized to β-tubulin or actin (n = 8) (right panel). ****: p < 0.0001. **(j)** PDK1 gene expression in p27^+/+^ (n = 8) and p27^-/-^ (n = 4) MEFs grown in full medium for 24 h from RNA-Seq data expressed as Count Per Million, normalized to p27^+/+^ levels. Graph shows means ± SEM. ****: p < 0.0001. **(k)** PDK1 mRNA level in p27^+/+^ and p27^-/-^ MEFs by RT-qPCR, normalized to p27^+/+^ levels. Graph shows means ± SEM from 5 independent experiments. **: p < 0.01. (**l**) Immunoblots for PDK1, p27 and actin in p27^+/+^ and p27^-/-^ MEFs (left panel). Graph shows means ± SEM of the quantification of PDK1 levels normalized to actin (n = 6) (right panel). *: p < 0.05. **(m)** Immunoblots for Lys18-lactylated Histone H3 (HH3-K18-La), p27 and actin in p27^+/+^ and p27^-/-^ MEFs (left panel). Graph shows means ± SEM of the quantification of HH3-K18-La levels normalized to actin (n = 6) (right panel). **: p < 0.01. **(n)** ATP quantification in p27^+/+^ and p27^-/-^ MEFs treated with either vehicle or Sodium Oxamate (50 mM) for 1 h. Graph shows means ± SEM from four independent experiments. *: p < 0.05; ***: p < 0.001.

Interestingly, Histone H3 (HH3) lactylation was described recently as an epigenetic mark that participates in the regulation of gene expression triggered by metabolic changes ^65^. Since lactate production is increased in p27^-/-^ MEFs, HH3 lactylation was evaluated by immunoblot. In p27^-/-^ cells, HH3 lactylation was significantly increased (Figure 5m) compared to p27^+/+^ cells, suggesting that lactate overproduction in p27^-/-^ cells leads to increased HH3 lactylation, which in turn could participate in modulating gene expression.

Most cancer cells exhibit a Warburg effect, accompanied by an increase in LDHA activity ^37,38,66^, as seen in p27^-/-^ MEFs. Elevated LDHA activity in cancers is associated with worst outcome ^67^, and LDHA inhibition with Sodium Oxamate (NaOxamate) has demonstrated some efficacy in cancers by decreasing ATP production in cancer cells ^68–70^. Sodium Oxamate treatment significantly decreased ATP production in p27^-/-^ cells compared to p27^+/+^ MEFs (Figure 5n). Thus, the loss of p27 in mouse fibroblasts appears to be sufficient to cause a Warburg effect and p27^-/-^ cells exhibit increased sensitivity to LDHA inhibition for ATP production, as reported in cancer cells ^37,38,66–70^.

### p27 controls glucose uptake

Cells lacking p27 appear to favor glycolysis and exhibit increased lactate production compared to wild-type cells. To gain further insight on the role of p27 in glucose metabolism, glucose levels in cells and culture media were measured in function of p27 status. p27^-/-^ cells had a significant decrease in glucose levels in their medium compared to fresh medium, suggesting elevated glucose uptake by these cells (Figure 6a). In contrast, intracellular glucose levels were similar in p27^+/+^ and p27^-/-^ MEFs (Figure 6b), probably reflecting the increased glycolytic capacity of p27^-/-^ cells. To explain the difference in glucose levels in culture media and determine whether p27^-/-^ cells exhibit increased glucose uptake, the expression of the ubiquitous glucose transporter GLUT1 was examined ^71^. RNA-Seq data, confirmed by RT-qPCR, revealed that GLUT1 expression is upregulated in p27^-/-^ MEFs compared to p27^+/+^ (Figure 6c, d). The increased expression of the GLUT1 protein was confirmed by immunoblot and immunostaining (Figure 6e, f).

**Figure 6:**
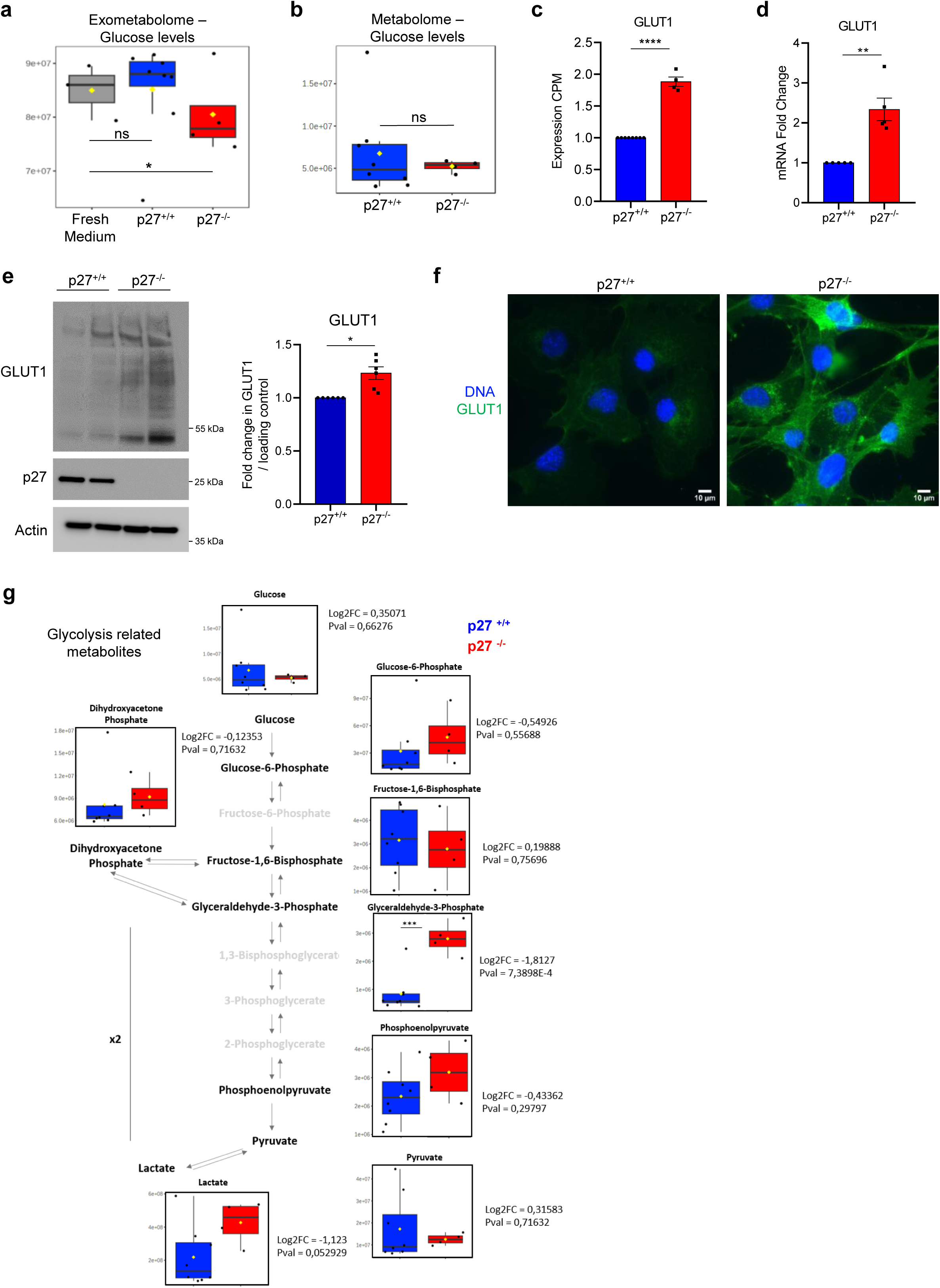
p27 controls glucose uptake. **(a)** Glucose quantification in fresh culture medium (grey) (n = 3), and media from p27^+/+^ (n = 8) and p27^-/-^(n = 4) MEFs grown for 24 h in full medium from exometabolome data. Data is presented as box plot, means are represented by yellow squares. ns: p > 0.05; *: p < 0.05. **(b)** Glucose quantification in p27^+/+^ (n = 8) and p27^-/-^ (n = 4) MEFs grown for 24 h in full medium from metabolome data. Data is presented as box plot, means are represented by yellow squares. ns: p > 0.05. **(c)** GLUT1 gene expression in p27^+/+^ (n = 8) and p27^-/-^ (n = 4) MEFs from RNA-Seq data expressed as Count Per Million, normalized to p27^+/+^ levels. Graph shows means ± SEM. ****: p < 0.0001**. (d)** GLUT1 mRNA levels in p27^+/+^ and p27^-/-^MEFs by RT-qPCR, normalized to p27^+/+^ levels. Graph shows means ± SEM from five independent experiments. **: p < 0.01. **(e)** Immunoblot for GLUT1, p27 and actin in p27^+/+^ and p27^-/-^ MEFs (left panel). Graph shows means ± SEM of the quantification of GLUT1 levels normalized to Actin (n = 6) (right panel). *: p < 0.05. **(f)** Representative images of GLUT1 immunostaining (green) in p27^+/+^ and p27^-/-^ MEFs. DNA was stained with H33342 (blue). **(g)** Graphical representation of glycolysis with metabolites from metabolome data in p27^+/+^ (n = 8) and p27^-/-^ (n = 4) MEFs grown for 24 h in full medium. Data is presented as box plots, means are represented by yellow squares. P values are indicated beside each graph.

To determine whether glycolytic flux was affected by p27 status, glycolysis metabolites levels were quantified by metabolomics experiments (Figure 6g). Although only G3P was significantly affected, all the glycolysis metabolites analyzed showed a similar trend and were elevated in p27^-/-^ MEFs compared to p27^+/+^ cells, supporting that p27 loss increases both glucose uptake and metabolism. These differences were abolished during glucose starvation and some metabolites (G3P, PEP, lactate) were even elevated in p27^+/+^ MEFs (Figure S2a), possibly due to their ability to increase autophagy flux during glucose deprivation ^19^. On the other hand, under amino acid starvation conditions, p27^-/-^ cells behaved largely like in full medium (Figure S2b). Taken together, these data indicate that p27 loss increases both glucose uptake and metabolism via the upregulation of GLUT1 and of glycolytic enzymes.

### Loss of p27 increases glutamine uptake and causes a glutamine dependency for mitochondrial respiration

In cancer cells, the Warburg effect is often coupled with a glutamine addiction, as they increase their use of glutamine as a carbon source to feed the TCA cycle ^49,72,73^. Since metabolomics studies indicate that glutamine levels are reduced in p27^-/-^ cells and medium compared to p27^+/+^ cells (Figure 3a-c), glutamine uptake and metabolism were investigated in more details. Glutamine level in the medium of p27^-/-^ cells was dramatically reduced compared to fresh medium and that of p27^+/+^ MEFs after 24 h of culture (Figure 7a). Similarly, intracellular glutamine levels in p27^-/-^ cells were significantly reduced compared to p27^+/+^ MEFs, suggesting that glutamine uptake and metabolism are increased in absence of p27. Indeed, expression levels of the glutamine transporter SLC1A5 was upregulated in p27^-/-^ cells compared to p27^+/+^ in RNA-Seq data, confirmed by RT-qPCR (Figure 7c-d) ^50^.

**Figure 7:**
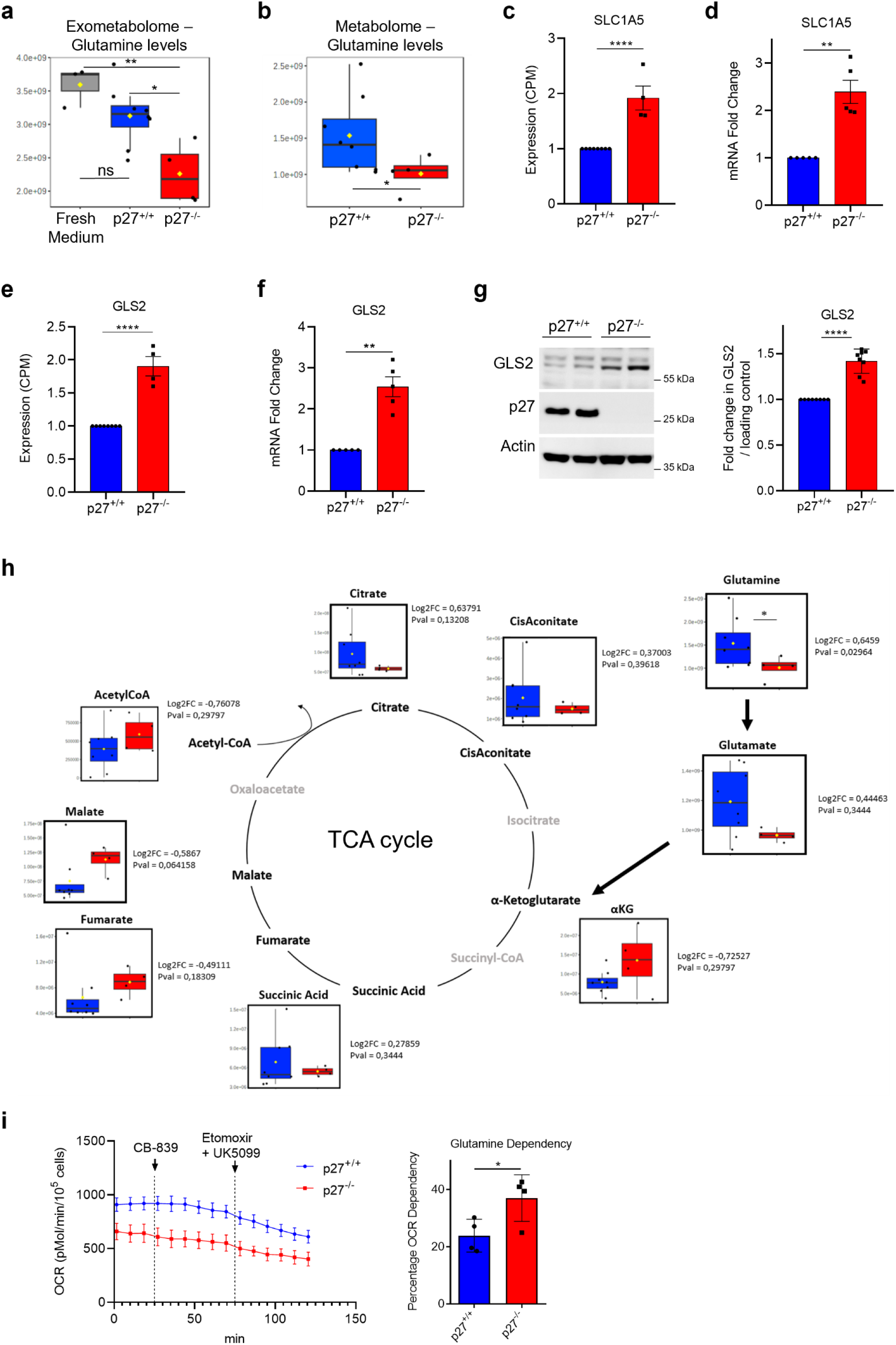
Loss of p27 increases glutamine uptake and causes a glutamine dependency for mitochondrial respiration. **(a)** Glutamine quantification in fresh culture medium (grey) (n = 3), and media from p27^+/+^ (n = 8) and p27^-/-^ (n = 4) MEFs grown for 24 h in full medium from exometabolome data. Data is presented as box plot, means are represented by yellow squares. ns: p > 0.05; *: p < 0.05; **: p < 0.01. **(b)** Glutamine quantification in p27^+/+^ (n = 8) and p27^-/-^ (n = 4) MEFs grown for 24 h in full medium from metabolome data. Data is presented as box plot, means are represented by yellow squares. *: p < 0.05. **(c)** SLC1A5 gene expression in p27^+/+^ (n = 8) and p27^-/-^ (n = 4) MEFs from RNA-Seq data expressed as Count Per Million, normalized to p27^+/+^ levels. Graph shows means ± SEM. ****: p < 0.0001. **(d)** SLC1A5 mRNA level in p27^+/+^ and p27^-/-^ MEFs by RT-qPCR, normalized to p27^+/+^ levels. Graph shows means ± SEM from five independent experiments. **: p < 0.01. **(e)** GLS2 gene expression in p27^+/+^ (n = 8) and p27^-/-^ (n = 4) MEFs from RNA-Seq data expressed as Count Per Million, normalized to p27^+/+^ levels. Graph shows means ± SEM. ****: p < 0.0001. **(f)** GLS2 mRNA level in p27^+/+^ and p27^-/-^ MEFs by RT-qPCR, normalized to p27^+/+^ levels. Graph shows means ± SEM from five independent experiments. **: p < 0.01. **(g)** Immunoblot for GLS2, p27 and actin in p27^+/+^ and p27^-/-^ MEFs (left panel). Graph shows means ± SEM of the quantification of GLS2 levels normalized to Actin (n = 8) (right panel). ****: p < 0.0001. (**h**) Graphical representation of TCA cycle related metabolites in p27^+/+^ (n = 8) and p27^-/-^ (n = 4) MEFs grown for 24 h in full medium from metabolome data. Data is presented as box plots, means are represented by yellow squares. P values are indicated beside each graph. **(i)** Glutamine dependency for mitochondrial respiration in p27^+/+^ and p27^-/-^ MEFs from Seahorse fuel dependency experiments (left panel). Oxygen consumption rate (OCR) was measured at various time points with consecutive injections of CB-839 (3 µM), and UK5099 (4 µM)/Etomoxir (2 µM), and normalized to cell number. From these experiments, the Glutamine dependency for mitochondrial respiration of p27^+/+^ and p27^-/-^ MEFs was calculated (right panel). Graphs show means ± SEM from four independent experiments. *: p < 0.05.

Once inside the cell, glutamine undergoes glutaminolysis before entering the TCA cycle in the form of α-Ketoglutarate (α-KG) ^74^. The first step of glutaminolysis is carried by glutaminase (GLS2) that transforms glutamine into glutamate. GLS2 levels were elevated at the mRNA and protein levels in p27^-/-^cells compared to p27^+/+^ MEFs (Figure 7e-g), suggesting that glutaminolysis is also enhanced in absence of p27. We then analyzed select TCA cycle metabolites by targeted metabolomics experiments, revealing that several of these metabolites (α-KG, fumarate and malate), all intermediates after the α-KG step that marks the point of entry in the TCA cycle fed by glutamine, had an elevated trend in p27^-/-^ MEFs compared to p27^+/+^ (Figure 7h). As anticipated, a similar trend was observed under glucose starvation, with elevated glutamine utilization to feed the TCA cycle (Figure S3a), which was abolished in absence of amino acids (Figure S3b), as expected. These results strongly suggest that p27^-/-^ MEFs use more glutamine as a substrate for the TCA cycle by increasing both uptake and glutaminolysis.

To test whether p27^-/-^ MEFs develop a glutamine addiction for mitochondrial respiration, as observed in cancer cells ^49^, we performed Seahorse Fuel Dependency experiments to calculate glutamine dependency in p27^+/+^ and p27^-/-^ MEFs, by sequentially injecting drugs inhibiting the use of the different substrates of the TCA cycle (Figure S3c) ^75^. These experiments revealed that glutamine dependency is markedly increased in p27^-/-^ MEFs cells compared to p27^+/+^ (Figure 7i), confirming that cells lacking p27 use more glutamine as a carbon source to feed the TCA cycle. Taken together, this data shows that p27 loss induces a reprogramming of cellular metabolism, with the induction of the Warburg effect, which is coupled to an increased utilization of glutamine for mitochondrial respiration.

### p27 silencing is sufficient to confer a glycolytic phenotype in untransformed human cells

The knockout of p27 causes a glycolytic phenotype in immortalized MEFs. We investigated whether the silencing of p27 was sufficient to cause a similar behavior in untransformed human cells. For this, p27 expression was silenced with siRNA for 72 h in hTert-immortalized human retinal pigment cells (hTert-RPE1). The silencing of p27 reduced OCR and increased ECAR in hTert-RPE1 cells compared to control siRNA transfected cells (Figure 8 a-b), albeit to a lower degree than in murine knockout fibroblasts, most likely due to incomplete loss of p27 expression in the former. The energy map of these cells indicated that p27 silencing drives metabolism from aerobic toward a glycolytic behavior (Figure 8c). Monitoring of the expression levels of glycolytic enzymes that were upregulated in p27^-/-^ MEFs (namely HK1, GAPDH, ENO1, ALDOA) showed similar changes upon p27 silencing in human cells (Figure 8d-g). Similarly, LDHA and LDHB expression were altered in p27 silenced hTert-RPE1 cells, as observed in murine fibroblast lacking p27 (Figure 8h, i). Thus, this data indicates that p27 silencing is sufficient to drive metabolism toward a glycolytic behavior, decreasing mitochondrial respiration and increasing lactate production in untransformed human cells.

**Figure 8:**
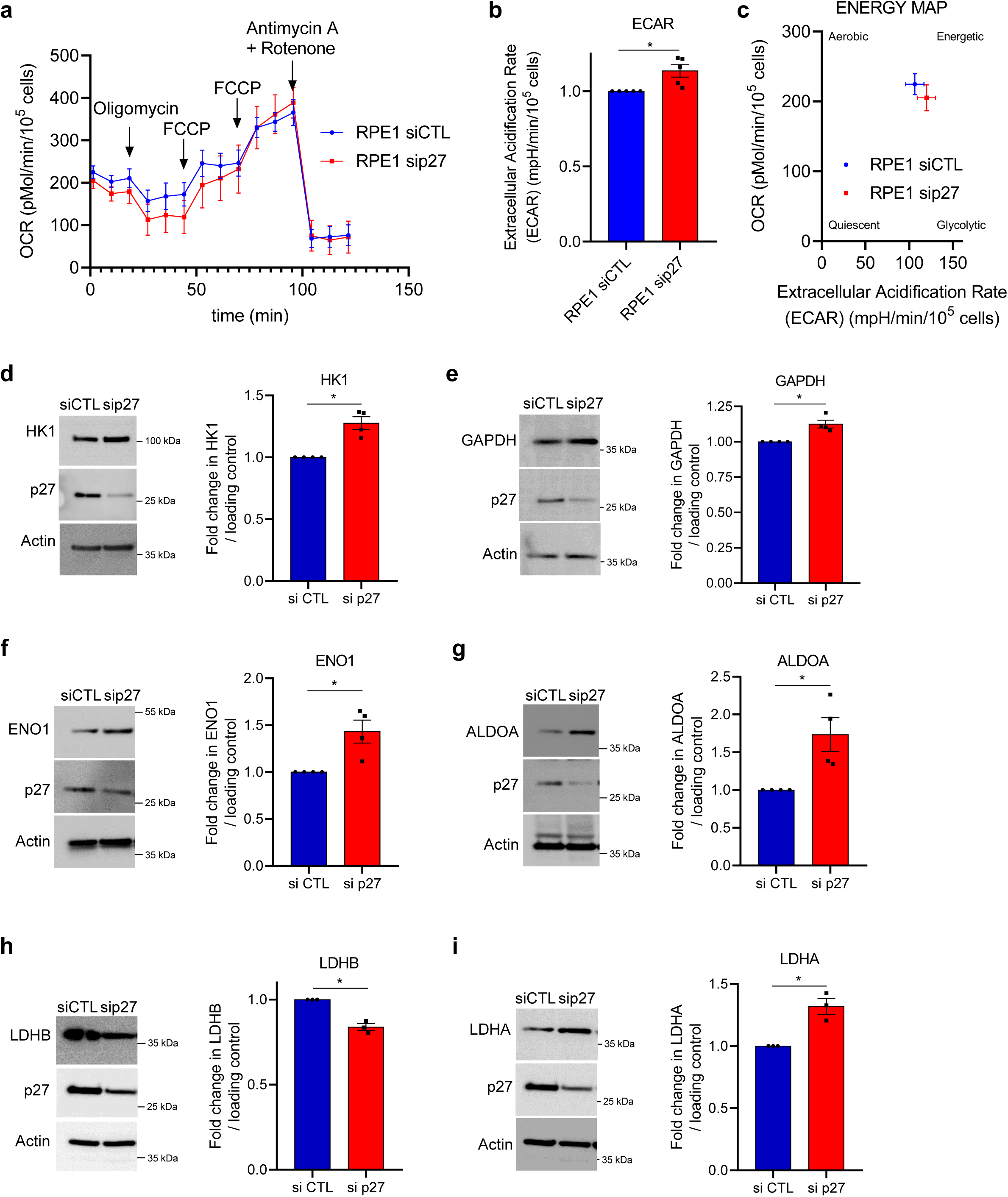
p27 silencing is sufficient to confer a glycolytic phenotype in RPE1-hTert human cells. **(a)** Oxygen consumption rate (OCR) was measured at various time points with consecutive injections of oligomycin (1 µM), FCCP (two injections at 2 µM and 6 µM), and Rotenone/antimycin A (1 µM each), and normalized to cell number. Graph shows the mean OCR curves of hTert-RPE1 transfected with control siRNA (siCTL) or p27 siRNA (sip27) (n = 5). **(b)** Extracellular acidification rate (ECAR) of p27 siRNA transfected hTert-RPE1 normalized to control siRNA condition (n = 5). Graph shows means ± SEM. *: p < 0.05. (**c**) Energy map of hTert-RPE1 transfected with control siRNA or p27 siRNA from Seahorse Mitostress experiments (n = 5). (**d**) Immunoblots for Hexokinase 1 (HK1), p27 and actin (loading control) in hTert-RPE1 transfected with control siRNA or p27 siRNA (left panel). Graph shows means ± SEM of the quantification of HK1 levels normalized to actin (n = 4) (right panel). *: p < 0.05. (**e**) Immunoblots for GAPDH, p27 and actin (loading control) in hTert-RPE1 transfected with control siRNA or p27 siRNA (left panel). Graph shows means ± SEM of the quantification of GAPDH levels normalized to actin (n = 4) (right panel). *: p < 0.05. (**f**) Immunoblots for Enolase 1 (ENO1), p27 and actin in hTert-RPE1 transfected with control siRNA or p27 siRNA (left panel). Graph shows means ± SEM of the quantification of ENO1 levels normalized to actin (n = 4) (right panel). *: p < 0.05. (**g**) Immunoblots for Aldolase A (ALDOA), p27 and actin in hTert-RPE1 transfected with control siRNA or p27 siRNA (left panel). Graph shows means ± SEM of the quantification of ALDOA levels normalized to actin (n = 4) (right panel). *: p < 0.05. (**h**) Immunoblots for Lactate Dehydrogenase B (LDHB), p27 and actin in hTert-RPE1 transfected with control siRNA or p27 siRNA (left panel). Graph shows means ± SEM of the quantification of LDHB levels normalized to actin (n = 3) (right panel). *: p < 0.05. (**i**) Immunoblots for Lactate Dehydrogenase A (LDHA), p27 and actin in hTert-RPE1 transfected with control siRNA or p27 siRNA (left panel). Graph shows means ± SEM of the quantification of LDHA levels normalized to actin (n = 3) (right panel). *: p < 0.05.

Altogether, our data suggest that the loss of p27 expression is sufficient to cause a metabolic reprogramming in fibroblasts and epithelial cells similar to that observed in cancer cells, with a Warburg effect coupled to a glutamine addiction.

## Discussion

To maintain cellular and tissue homeostasis, a crosstalk between the metabolic and cell cycle machineries is essential to sustain proliferation and growth, as cell cycle progression is strongly dependent on energetic output ^1–4^. While it has been firmly established that metabolic pathways can control cell cycle progression, increasing evidence point to reciprocal interactions where cell cycle regulators also control metabolic pathways to modulate metabolic output ^1–4^. Indeed, the role of the cyclin/CDK complexes/Rb/E2F axis has been extensively studied ^1–6,76^. However, the importance of CKIs in this crosstalk is still largely unclear. Nevertheless, p21 and p15^INK4B^ have been involved in modulating glycolysis ^7,77^, while fatty acid metabolism was reported to be regulated by p21 and p16 ^8,9^. Similarly, a number of studies have investigated the role of p27 in the control of autophagy induction in response to metabolic stress ^19–22^, but the importance of p27 in modulating energetic metabolism remains largely unknown. In this study, using bioenergetics, metabolomics and transcriptomics approaches, we found that p27 plays a role in energy metabolism by regulating glucose and glutamine metabolism. Indeed, our data indicate that loss of p27 expression is sufficient to induce extensive metabolic reprogramming, causing a Warburg effect and a glutamine addiction.

These findings are particularly relevant in the context of cancer, in which p27 is frequently inactivated either by increased degradation or mislocalization in the cytoplasm, and more rarely through mutations ^10,12,14,15^. Moreover, p27 mislocalization appears to occur early during tumorigenesis as a secondary event following activation of oncogenic pathways such as K-Ras, BCR-Abl or PI-3K/Akt signaling ^12,28,78,79^. For instance, cytoplasmic localization of p27 is induced by K-Ras activation in pre-neoplastic lesions in the pancreas, or by *Helicobacter pylori* infection of gastric epithelial cells via PI-3K/Akt activation ^28,78^. Thus, in these early stages of cancer development, p27 inactivation through degradation or mislocalization would both remove a cell cycle brake, enabling unlimited proliferation, and cause a metabolic reprogramming leading to a Warburg effect and increased glutamine metabolism, thereby providing the fuel and energy required for this deregulated proliferation.

Overall, ATP production is similar in p27^+/+^ and p27^-/-^ MEFs, and increased glycolytic flux seems to compensate for the decreased production of mitochondrial ATP. p27^-/-^ MEFs are more sensitive to apoptosis induced by glucose deprivation and this was associated with their decreased ability to perform autophagy ^19,20^. However, in light of our data, it appears likely that the increase dependency of p27^-/-^ cells on glycolysis may also participate in this increased susceptibility to cell death in absence of glucose. In the context of cancer, it would be interesting to test the response of p27^+/+^ and p27^-/-^ cells to drugs targeting glycolysis, such as 2-DG, or lactate production, such as Sodium Oxamate, already used in the clinic ^68–70,80^. Our data also indicate that cells lacking p27 are more dependent on glutamine for ATP generation than their wild-type counterparts and interfering with glutamine metabolism, either its transport or glutaminolysis, possibly in combination with glycolysis inhibitors, could be an efficient strategy to target these cells ^81^.

Our data indicate that loss of p27 expression triggers a metabolic reprogramming mediated at least in part via transcriptional changes that lead to increased expression of all ten glycolytic enzymes and the glucose transporter GLUT1, of LDHA that favors lactate production, and the glutamine transporter SLC1A5 and the glutaminase GLS2. p27 was reported to regulate transcription by interacting with several transcriptional corepressors, such as mSin3a, and HDAC1, −4 and −5, and transcription factors ^11,23–27^, and p27 could be directly involved in the repression of these genes in homeostatic conditions. In addition, although it is a secondary event caused by lactate overproduction in p27^-/-^ MEFs, histone lactylation could contribute in reinforcing the gene expression changes triggered by loss of p27 expression. Indeed, we observed elevated levels of H3K18la, known to modify gene expression linked to metabolism ^82^, and another histone lactylation, H4K12la, was reported to increase expression of PKM2 and LDHA ^83^, leading to a positive feedback on lactate production and participating in establishing the Warburg effect.

Together, these findings reveal that loss of p27 expression is sufficient to cause metabolic reprogramming, at least in part through the deregulation of genes involved in glycolysis and glutamine metabolism, and underscore the fact that in a single event, p27 inactivation causes the loss of cell cycle control and triggers metabolic changes that can sustain deregulated proliferation. These data suggest that inhibiting glucose and/or glutamine metabolism in cancer cells in which p27 has been inactivated could be an efficient strategy to target these cells.

## Materials and Methods

### Reagents and antibodies

Control siRNA (sc-44232) and p27 siRNA (sc-29429) were purchased from Santa Cruz Biotechnology and transfected with Interferin (Polyplus transfection).

Mouse anti-p27 (SX53G8.5, sc-53871; Immunofluorescence (IF) 1/100, Western Blot (WB) 1/1000), Aldolase A (sc-377058; WB 1/1000), Enolase 1 (sc-271384; WB 1/1000) antibodies were purchased from Santa Cruz Biotechnologies. Mouse anti-p27 (BD, 610242; WB 1/500) was purchased from BD Transduction Laboratories. Rabbit anti-Hexokinase 1 (C35C4; WB 1/1000), GAPDH (D16H11; WB 1/1000), PKM1/2 (C103A3; WB 1/1000), LDHA (2012S, WB 1/1000), GLUT1 (E4S6I; WB 1/1000; IF 1/100), PDK1 (3820S; WB 1/1000), β-Tubulin (2128S, WB 1/1000) were purchased from Cell Signalling Technology. Rabbit anti-LDHB (A18096; WB 1/1000), GLS2 (A16029; WB 1/1000) were purchased from ABclonal. Rabbit anti-L-lactyl-Histone H3 (Lys18) (22H2L3; WB 1/1000) was purchase from PTM BIO.

All secondary antibodies were purchased from Jackson ImmunoResearch Laboratory.

### Cell lines, culture and transfection

hTERT RPE-1 cells were kindly provided by Dr. Olivier Calvayrac (Centre de Recherche en Cancérologie de Toulouse, France).

Primary mouse embryonic fibroblasts (MEFs) were prepared as described previously from p27^+/+^ and p27^-/-^ embryos ^84,85^. Several lines of p27^+/+^ and p27^-/-^ MEFs were immortalized with retroviruses encoding the human papillomavirus E6 protein and selected with hygromycin (250 µg/mL), as described previously^20^. Cells were grown in DMEM (10-013-CV, Corning) 4.5 g/L glucose supplemented with 10% fetal bovine serum (FBS) (F7524, Sigma), 0.1 mM non-essential amino acids (M7145, Sigma) and 2 μg/mL penicillin–streptomycin (15070063, Corning) at 37 °C, 5% CO2.

All cell lines were tested regularly for mycoplasma contamination using MycoAlert® Mycoplasma Detection Kit (LT07-318, Lonza).

For amino acid starvation experiments, cells were washed three times with PBS and grown with DMEM without amino acids (D9800, USBiological) complemented to 4.5 g/L D-glucose (88270, Sigma), 0.1 mM sodium pyruvate (11360039, Gibco), 2 μg/mL penicillin/streptomycin and 10% dialyzed FBS for 24 h. For glucose starvation experiments, cells were washed three times with PBS and then cultivated with glucose-free DMEM (D5030, Sigma-Aldrich) supplemented with 0.1 mM nonessential amino acids, 2 μg/ml penicillin–streptomycin, 2 mM L-glutamine (G7513, Sigma) and 10% dialyzed FBS for 24 h. For all starvation experiments, FBS was dialyzed against PBS in dialysis tubing with a 3,500 MW cut-off (Spectrum Labs, 132111) following the manufacturer’s instructions.

For siRNA Transfection, hTERT RPE-1 cells were transfected twice with control siRNA or p27 siRNA for 72 h (20 nM for 24 h followed by another transfection with 20 nM for 48 h) using Interferin (101000016, Polyplus) according to the manufacturer’s instructions.

### Immunoblotting

All immunoblots were performed on at least to distinct MEF lines per genotype.

Cells were lysed in lysis buffer (HEPES 50 mM pH 7.5, NaCl 150 mM, EDTA 1 mM, EGTA 2.5 mM, NP-40 1%, Tween 20 0.1%, glycerol 10%, supplemented with dithiothreitol 1 mM, phosphatase inhibitors (b-glycerophosphate, NaF and sodium orthovanadate at 10 mM each) and protease inhibitors (Aprotinin, Bestatin, Leupeptin and Pepstatin A, each at 10 µg/mL). Cell lysates were sonicated for 10 s and the protein concentration was determined by Bradford assay. Proteins were resolved on 4-20% gradient SDS-PAGE gels, transferred to PVDF membrane (Immobilon-P, Millipore) and blocked for 1 h in PBS-T (PBS 0.1% Tween 20) containing 5% non-fat dry milk. Membranes were incubated with the indicated primary antibodies overnight at 4 °C under agitation in PBS-T 5% BSA, washed three times in PBS-T and probed with HRP-conjugated secondary antibodies (1/10000 dilution) for 4 h at room temperature. Signals were visualized with chemiluminescence detection reagent (1705060S, Bio-Rad) using a Fusion Solo S (Vilber) or a qTOUCH (RWD Life Science Co.) digital acquisition system. Immunoblot signal intensity was evaluated using the ImageJ software and normalized to corresponding loading control intensity (β-Actin or β-Tubulin).

### Immunofluorescence

MEFs were seeded on coverslips, grown to 50-60% confluence and fixed with 4% PFA for 10 min at room temperature. Coverslips were washed three times for 5 min with PBS and blocked for 20 min with blocking solution (PBS, 3% BSA, 0.05% Tween-20 and 0.08% sodium azide) at room temperature. Cells were permeabilized for 3 min in PBS 0.2% Triton-X100 and washed 3 times in PBS. Then, coverslips were incubated with primary antibodies in blocking solution for 1 h at 37 °C, washed three times for 5 min in PBS, and incubated with Cy2-conjugated secondary antibodies for 30 min at 37 °C. Coverslips were washed once with PBS containing 0.1 μg/mL Hoechst H33342 for 5 min and twice in PBS before being mounted on glass slides using gelvatol (10% polyvinyl alcohol (w/v), 20% glycerol (v/v), 70 mM Tris pH 8). Image acquisition was performed on a Nikon 90i Eclipse microscope and a DS-Qi2 HQ camera using the NIS Element BR software.

### Seahorse experiments

XF24 cell culture plates (Agilent Technologies) were used to measure Oxygen Consumption Rate (OCR) and Extra Cellular Acidification Rate (ECAR). XF24 wells were coated with 0.2% gelatin for 30 min at 37 °C, then gelatin excess was removed and the plates were dried overnight at 37 °C. The sensor cartridge was hydrated overnight with calibration buffer supplied by Seahorse Biosciences. All Seahorse experiments were performed on at least to distinct MEF lines per genotype.

#### Seahorse Mito Stress experiments

1 x 10^4^ cells were seeded per well for 24 h, the medium was then replaced by XF base minimal DMEM medium containing 11 mM glucose, 1 mM pyruvate, and 2 mM glutamine. Cells were placed in a CO_2_-free incubator at 37 °C for 1 h and the baseline OCR and ECAR were measured. Then, oligomycin (1 µM; O013, TOKU-E), an Electron Transport Chain (ETC) complex V inhibitor, was added and the resulting OCR was used to derive ATP-linked OCR and proton leak respiration. Carbonyl cyanide-p-trifluoromethoxy-phenyl-hydrazon (FCCP; C2579, Sigma), a protonophore causing the ETC to function at its maximal rate, was added twice (2 µM and 6 µM). The maximal OCR was derived by subtracting non-mitochondrial respiration from the FCCP rate. Finally, Antimycin A (1 µM; A8674, Sigma) and rotenone (1 µM; R8875, Sigma), ETC complex III and I inhibitors, respectively, were added to shut down the ETC function to determine the non-mitochondrial OCR. The OCR and ECAR dataset from Seahorse Mito Stress experiments were used to quantify ATP production and glycolytic/mitochondrial ATP using the Agilent Seahorse XF ATP Real-Time rate method.

#### Seahorse Glycolysis Stress experiments

Cell media of XF24 plates (1 x 10^4^ cells/well) was replaced with XF base minimal DMEM medium containing 2 mM glutamine and cells were placed in a CO_2_-free incubator at 37 °C for 1 h. At the end of this incubation, the baseline ECAR was measured. Glucose (10 mM) was added to derive glycolysis linked ECAR, followed by Oligomycin (1 µM) to derive the Glycolytic Capacity of cells and 2-deoxy-D-glucose (2-DG; 50 mM, T6742, TargetMol) to derive non-glycolytic ECAR. The ECAR linked to glucose consumption by the cells (glycolytic ECAR) was calculated by subtracting non-glycolytic ECAR from each ECAR measure.

#### Seahorse Fuel dependency experiments

cells were treated as for Seahorse Mito Stress experiments to measure baseline OCR. Then, CB-839 (3 µM; PHR1084, Selleckchem), a glutaminase inhibitor, was added first followed by UK-5099 (4 µM; 5.04817, Sigma), a Pyruvate carrier inhibitor, and Etomoxir (2 µM; HY-50202A; MedChemExpress), a long-chain fatty acid oxidation inhibitor and the resulting OCR was used to calculate glutamine dependency using the following equation: Dependency (%) = ((Baseline – Glutaminase Inhibitor) / (Baseline – All inhibitors)) x 100.

For all Seahorse experiments, OCR and ECAR data were normalized to the cell number in each well of the XF24 cell plates. For this, at the end of each Seahorse experiment, cells were stained with 0.1 μg/mL Hoechst H33342, the entire wells were imaged using an EVOS M7000 microscope (Thermo Fisher Scientific) using a 4X objective and nuclei were counted using the Fiji plugin Stardist.

### Targeted LC–MS metabolomics cell preparation, analyses and data treatment

Two lines of p27^+/+^ and one line of p27^-/-^ MEFs were seeded in 60 mm plates until 40-50% confluency. The next day, cell medium was replaced with either full medium, glucose starvation medium, or amino acid starvation medium, as described above. After 24 h, cells were washed three times with chilled PBS and frozen in liquid nitrogen before metabolite extraction. Four independent experiments were performed for each line and culture condition.

Metabolites were extracted using a solution consisting of 50% methanol, 30% acetonitrile (ACN), and 20% water. The volume of extraction solvent was normalized to cell number (1 mL per 1×10[ cells). Following the addition of the extraction solution, samples were vortexed for 5 min at 4 °C and centrifuged at 16,000 × g for 15 min at 4 °C. The resulting supernatants were collected and stored at−80 °C until analyzed.

Liquid chromatography–mass spectrometry (LC/MS) was performed using a QExactive Plus Orbitrap mass spectrometer equipped with an Ion Max source and HESI II probe, coupled to a Dionex UltiMate 3000 UPLC system (Thermo Fisher Scientific). A 5 µL aliquot of each sample was injected onto a ZIC-pHILIC column (150 × 2.1 mm, 5 µm particle size) with a corresponding guard column (20 × 2.1 mm, 5 µm; Millipore) for separation.

The mobile phase consisted of buffer A (20 mM ammonium carbonate, 0.1% ammonium hydroxide, pH 9.2) and buffer B (ACN). The chromatographic gradient was run at 0.2 µL/min as follows: 0–20 min, linear decrease from 80% to 20% buffer B; 20–20.5 min, linear increase back to 80% buffer B; 20.5–28 min, held at 80% buffer B.

The mass spectrometer operated in full scan, polarity switching mode. Settings included a spray voltage of 2.5 kV, a heated capillary temperature of 320 °C, sheath gas flow of 20 units, auxiliary gas flow of 5 units, and sweep gas flow of 0 units. Metabolites were detected over a mass range of 75–1000 m/z at a resolution of 35,000 (at 200 m/z), with an automatic gain control (AGC) target of 1×10[ and a maximum injection time of 250 ms. Lock masses were used to maintain mass accuracy within 5 ppm.

Data acquisition was performed using Thermo Xcalibur software. Metabolites and isotopologues were quantified by peak area using Thermo TraceFinder software, based on their exact mass (as singly charged ions) and known retention times.

Enrichment analyses were carried out using MetaboAnalyst 5.0 and the metabolite concentrations were normalized by sum and scaled by mean-centering method. Volcano plots were generated from data of metabolome, exometabolome or culture medium alone (without cells) by combining Fold Change (FC) analysis and t-tests into one single graph: Pval threshold = 0.05 (FDR); Fold Change threshold = 1.5. Heatmaps were generated using only metabolites statistically different between p27^+/+^ and p27^-/-^ MEFs derived from Volcano plots of metabolome or exometabolome. Box plots represent the original concentration of each metabolite, associated with log2(FC) and Pval derived from Volcano plots using normalized data. Box plot comparing culture medium alone and exometabolome were generated comparing data from culture medium alone versus p27^+/+^ or p27^-/-^ exometabolome by combining Fold Change (FC) Analysis and t-tests, associated in the same graph. Principal Component Analyses were performed using PERMANOVA based on Euclidean distance of normalized concentration of metabolites.

### Flow Cytometry

p27^+/+^ or p27^-/-^ MEFs were collected by trypsinization and resuspended in PBS. Cells were stained with 100 nM MitoTracker Green (MTG; M7514, Sigma) for 20 min at 37 °C. After washes, cells were incubated with anti-annexin V antibody (BD Horizon™ BV421 Annexin V; 1/100). After removing debris, the mitochondrial mass (MTG staining) was evaluated in at least 5 x 10^3^ live cells (Annexin V negative cells). Data was analyzed with the FlowJo v10 software.

### Reverse transcription (RT) and quantitative polymerase chain reaction (qPCR)

Extraction of total RNA from cultured cells and RT were performed as described previously ^86^. qPCR was performed using ONEGreen Fast qPCR Premix (Ozyme) on a CFX Opus 96 Real Time PCR system (BioRad) according to the manufacturer’s instructions. Results were analyzed using the CFX Manager Software (BioRad) using the 2-ΔΔCt method. The mRNA levels were normalized to β-microglobulin (B2m). Primers for qPCR were synthesized by Eurofins Genomics (Ebersberg, Germany).

#### RT-qPCR primers

Sequences of mouse primers used in qPCR experiments. FW: Forward, RV: Reverse.

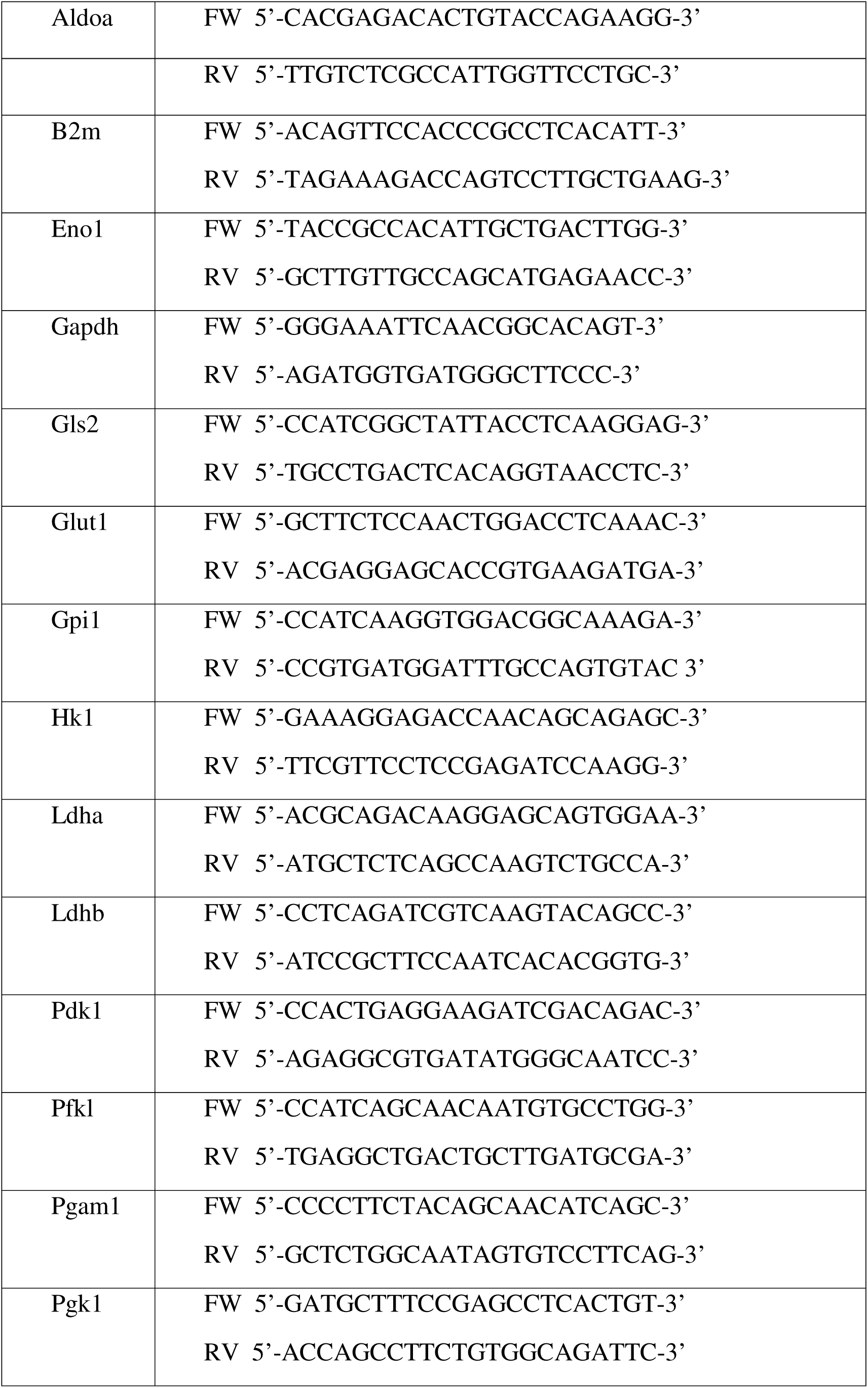
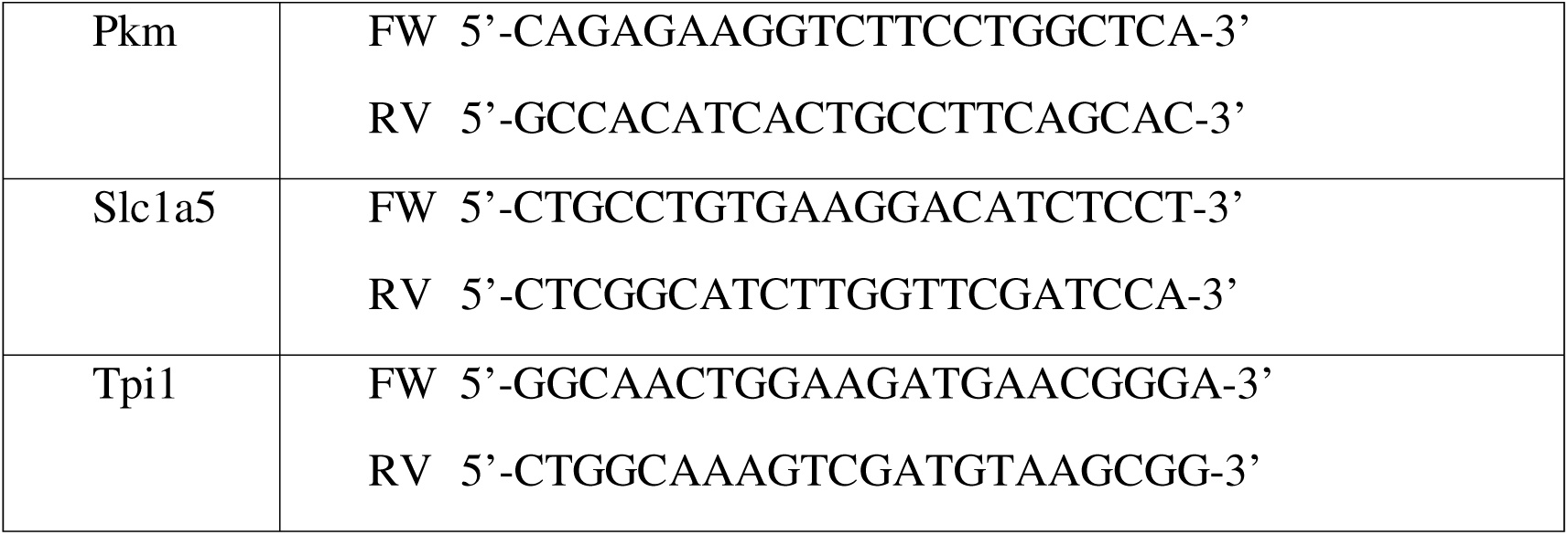

### ATP quantification

Cellular ATP was quantified using Firefly Luciferase ATP Assay kit (Cat # 28854, Cell Signalling Technology). Briefly, 4 x 10^3^ cells were seeded per well in 96 well plates. After 24 h, cells were treated with a mix of Oligomycin A (10 µM) and Antimycine A (10 µM) (OAA), or 2-DG (50 mM), or with a mix of OAA and 2-DG or vehicle for 1 h. ATP was quantified according to Firefly Luciferase ATP Assay kit instructions using a ThermoScientifc Luminoskan^TM^ luminometer. Luminescence intensity was normalized to cell number as described in the Seahorse experiments section. The luminescence intensity of the cells treated with of OAA + 2-DG represents the residual ATP and was subtracted from each condition. For ATP quantification during LDHA inhibition, cells were pre-treated with 50 mM Sodium Oxamate (50 mM) (O2751, Sigma Aldrich).

### RNA extraction, RNA-Seq and transcriptomics analyses

Two lines of p27^+/+^ (n = 8) and one line of p27^-/-^ (n = 4) grown in full medium for 24 h were used for transcriptomics analyses. Cells were lysed in TriReagent (T9424, Sigma Aldrich) and RNA was extracted using Direct-zol RNA Miniprep Plus Kits (Zymo Research) according to the manufacturer’s protocol. RNA integrity was verified on 0.8% agarose gel and quantified using a Nanodrop spectrophotometer. cDNA was synthetized using SuperScript IV Reverse Transcriptase (Thermo Fisher Scientific) according to the manufacturer’s protocol. RNA-Seq samples were sequenced using Illumina Novaseq (paired-end, 2 x 150 bp long reads, 10M reads/sample). The quality of each raw sequencing file (fastq) was verified with FastQC (https://www.bioinformatics.babraham.ac.uk/projects/fastqc/). Fastq were aligned the reference mouse genome (GRCm39) and counted on the GENCODE annotation from database (gtf GRCm39) using STAR aligner (STAR_2.5.4a) ^87^. Then the raw count table was cleaned using Bioconductor HTSfilter package with s.min = 1, s.max = 50 parameters (HTSFilter_1.44.0) ^88^. Differential analysis (normalization and generation of log2FC and adjusted p-value) was applied independently, for each p27^+/+^ against p27^-/-^using Bioconductor DESeq2 (DESeq2_1.44.0) ^89,90^. Genes were considered differentially expressed when −0.6 < log2(FC) < 0.6 and p < 0.05. We used principal component analysis (PCA) to cluster samples based on their normalized expression levels. RNA-Seq data have been submitted on the GEO database under accession number GSEXXX.

Gene Ontology was performed using ShinyGO 0.81, based on the 10 most enriched clusters of up- or down-regulated genes in p27^-/-^ compared to p27^+/+^ MEFs. CDKN1B coding sequence was visualized using Integrative Genomics Viewer based on p27^+/+^ and p27^-/-^ reads.

### Statistical analyses

Statistical analyses were performed using the Graphpad Prism 8.0 software. All statistical analyses are based on at least three independent experiments. Comparisons between two groups were performed using the unpaired t-test with Welch’s correction. Differences between three groups were performed using Two-way ANOVA tests followed by Sidak’s multiple comparisons when only genotypes are compared with each other, or followed by Tukey’s multiples comparisons when genotypes and treatments are compared with each other. Data are presented as means ± SEM. Symbols used are: ns: p > 0.05; *: p < 0.05; **: p < 0.01; ***: p < 0.001; ****: p < 0.0001.

Statistical analyses of metabolomics data are described in the section Targeted LC–MS metabolomics cell preparation, analyses and data treatment. Statistical analyses of transcriptomics data are described in the section RNA extraction, RNA-Seq and transcriptomics analyses.

## Author contributions

LR, CD, EM, AB and IN performed experiments. LR, AB, AH, JC and J-ES designed experiments. LR, MA, CD, AH and AB analyzed the data. LR, AH and AB wrote the paper with contributions from all authors.

## Acknowledgements

LR was supported by a Studentship from the Fondation pour la Recherche Médicale. EM was supported by a PhD Studentship from the Ministère de l’Enseignement Supérieur et de la Recherche. This work was supported by an FRM Equipes grant (EQU202403018042) from the Fondation pour la Recherche Médicale to AB.

---

**Figure S1:**
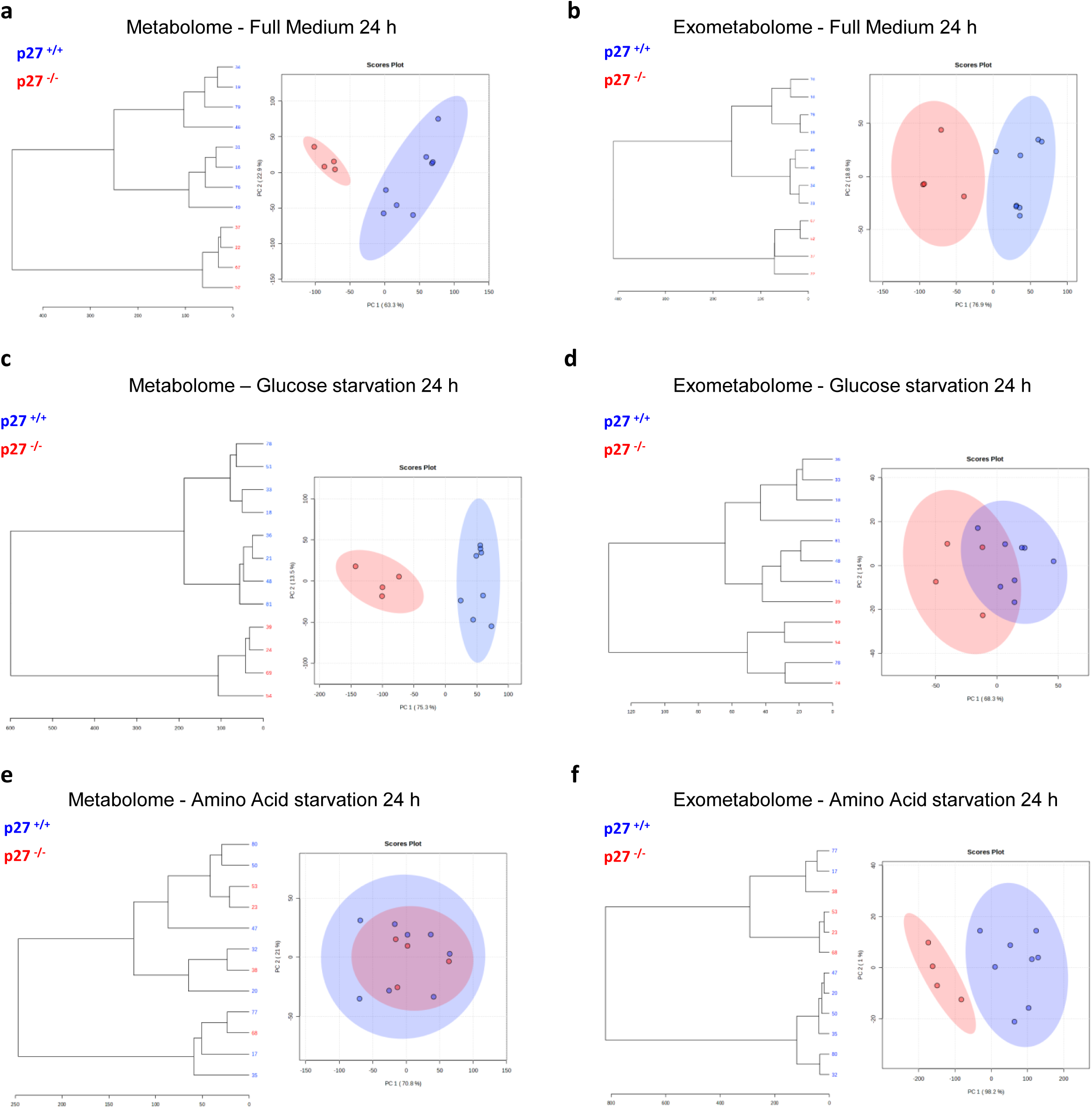
p27 impacts global metabolism and exometabolism during normal condition or metabolic stress. Dendrogram (left panel) and principal component analysis (right panel) of cellular metabolites (a) and exometabolites (b) in p27^+/+^ (n = 8) and p27^-/-^ (n = 4) MEFs after 24 h of culture in full medium **(a, b)**, or 24 h of culture in absence of glucose **(c, d)**, or 24 h of culture in absence of amino acids **(e, f)**.

**Figure S2:**
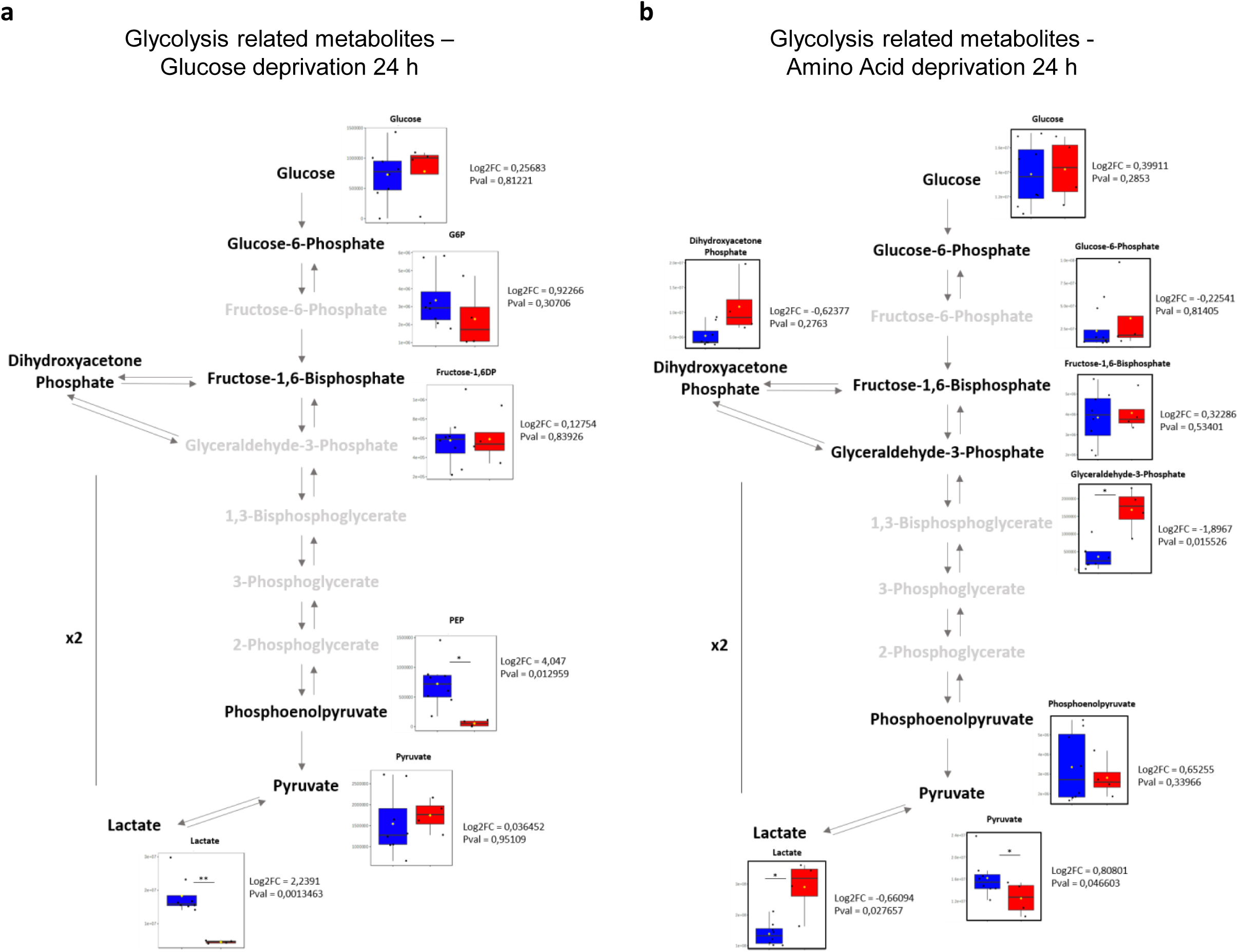
Analysis of glycolysis metabolites in p27^+/+^ and p27^-/-^ MEFs in glucose or amino acid starvation conditions. **(a, b)** Graphical representation of glycolysis showing cell metabolites analyzed in the metabolome of p27^+/+^ (n = 8) and p27^-/-^ (n = 4) MEFs after 24 h in glucose starvation conditions (a) or after 24 h in amino acid starvation conditions. Data is presented as box plot, means are represented by yellow squares. ns: p > 0.05; *: p < 0.05; **: p < 0.01. p values are indicated beside each box plot.

**Figure S3:**
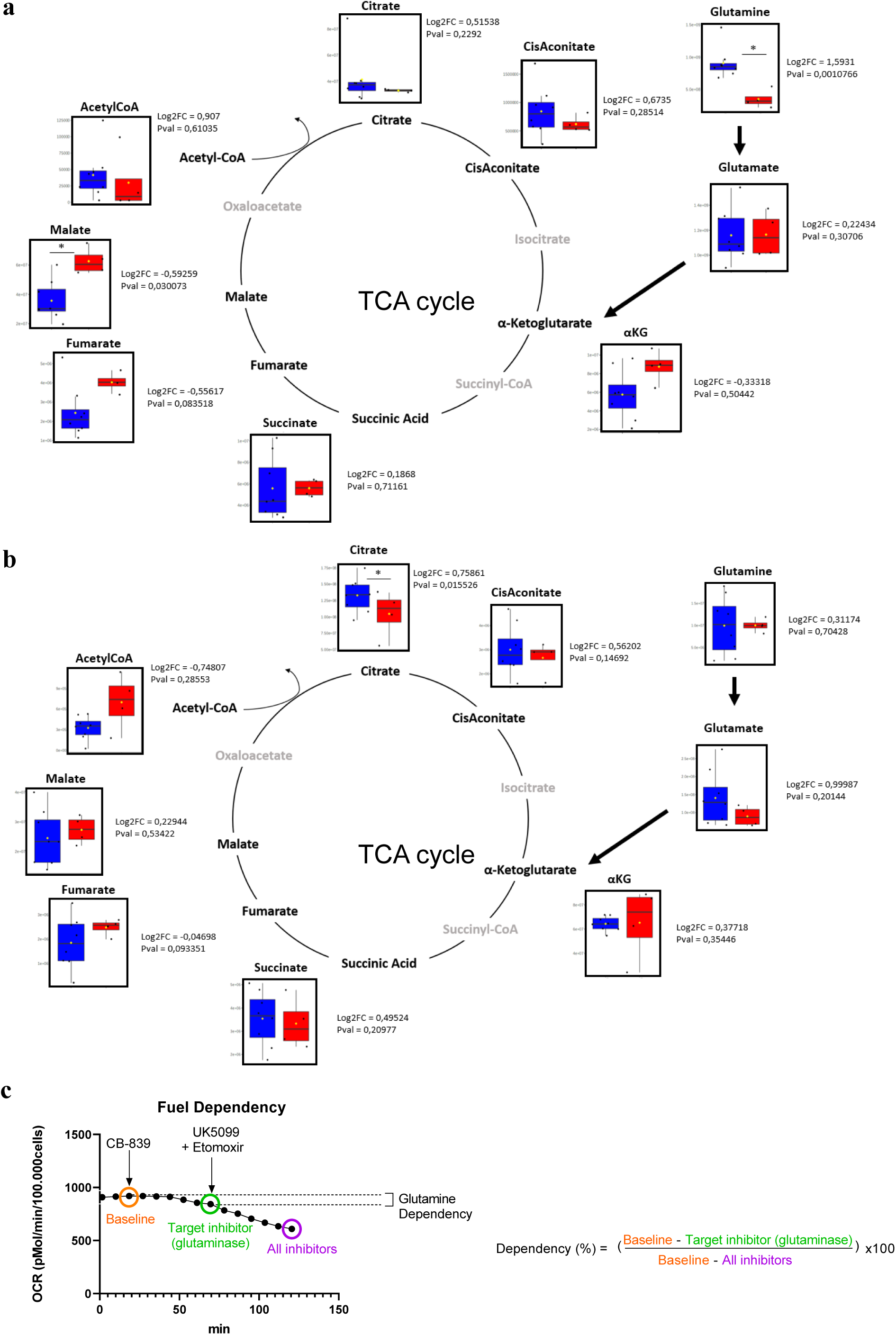
Analysis of TCA cycle metabolites in p27^+/+^ and p27^-/-^ MEFs under glucose or amino acid starvation. **(a, b)** Graphical representation of TCA cycle-related metabolites analyzed in the metabolome of p27^+/+^ (n = 8) and p27^-/-^ (n = 4) MEFs after 24 h in glucose starvation conditions (a) or after 24 h in amino acids starvation conditions (b). Data is presented as box plot, means are represented by yellow squares. ns: p > 0.05; *: p < 0.05. **(c)** Oxygen consumption rate (OCR) curve showing the calculation used to determine glutamine dependency for mitochondrial respiration in Seahorse fuel dependency experiments. Measurements were performed after injections of CB-839 (Glutaminase inhibitor; 3 µM), and UK5099 (inhibits pyruvate entry to mitochondria; 4µM) / Etomoxir (inhibits long-fatty acid entry to mitochondria; 2 µM)

